# Evolution of the bread wheat D-subgenome and enriching it with diversity from *Aegilops tauschii*

**DOI:** 10.1101/2021.01.31.428788

**Authors:** Kumar Gaurav, Sanu Arora, Paula Silva, Javier Sánchez-Martín, Richard Horsnell, Liangliang Gao, Gurcharn S. Brar, Victoria Widrig, Jon Raupp, Narinder Singh, Shuangye Wu, Sandip M. Kale, Catherine Chinoy, Paul Nicholson, Jesús Quiroz-Chávez, James Simmonds, Sadiye Hayta, Mark A. Smedley, Wendy Harwood, Suzi Pearce, David Gilbert, Ngonidzashe Kangara, Catherine Gardener, Macarena Forner-Martínez, Jiaqian Liu, Guotai Yu, Scott Boden, Attilio Pascucci, Sreya Ghosh, Amber N. Hafeez, Tom O’Hara, Joshua Waites, Jitender Cheema, Burkhard Steuernagel, Mehran Patpour, Annemarie Fejer Justesen, Shuyu Liu, Jackie C. Rudd, Raz Avni, Amir Sharon, Barbara Steiner, Rizky Pasthika Kirana, Hermann Buerstmayr, Ali A. Mehrabi, Firuza Y. Nasyrova, Noam Chayut, Oadi Matny, Brian J. Steffenson, Nitika Sandhu, Parveen Chhuneja, Evans Lagudah, Ahmed F. Elkot, Simon Tyrrell, Xingdong Bian, Robert P. Davey, Martin Simonsen, Leif Schauser, Vijay K. Tiwari, H. Randy Kutcher, Pierre Hucl, Aili Li, Deng-Cai Liu, Long Mao, Steven Xu, Gina Brown-Guedira, Justin Faris, Jan Dvorak, Ming-Cheng Luo, Ksenia Krasileva, Thomas Lux, Susanne Artmeier, Klaus F. X. Mayer, Cristobal Uauy, Martin Mascher, Alison R. Bentley, Beat Keller, Jesse Poland, Brande B. H. Wulff

## Abstract

*Aegilops tauschii,* the diploid wild progenitor of the D-subgenome of bread wheat, constitutes a reservoir of genetic diversity for improving bread wheat performance and environmental resilience. To better define and understand this diversity, we sequenced 242 *Ae. tauschii* accessions and compared them to the wheat D-subgenome. We characterized a rare, geographically-restricted lineage of *Ae. tauschii* and discovered that it contributed to the wheat D-subgenome, thereby elucidating the origin of bread wheat from at least two independent hybridizations. We then used *k*-mer-based association mapping to identify discrete genomic regions with candidate genes for disease and pest resistance and demonstrated their functional transfer into wheat by transgenesis and wide crossing, including the generation of a library of ‘synthetic’ hexaploids incorporating diverse *Ae. tauschii* genomes. This pipeline permits rapid trait discovery in the diploid ancestor through to functional genetic validation in a hexaploid background amenable to breeding.

## Main

The success of bread wheat (*Triticum aestivum*) as a major worldwide crop is underpinned by its adaptability to diverse environments, high grain yield and nutritional content^1^. With the combined challenge of population expansion and hotter, less favorable climates, wheat yields must be sustainably increased to ensure global food security. The rich reservoir of genetic diversity amongst the wild relatives of wheat provides a means to improve productivity^1, 2^. Maximizing the genetic potential of wheat requires a deep understanding of the structure and function of its genome, including its relationship with its wild progenitor species.

The evolution of bread wheat from its wild relatives is typically depicted as two sequential interspecific hybridization and genome duplication events leading to the genesis of the allohexaploid bread wheat genome^2, 3^. The first hybridization between *T*. *urartu* (AA) and a presumed extinct diploid (BB) species formed tetraploid emmer wheat, *T*. *turgidum* (AABB), ∼0.5 million years ago^4^. The gradual process of domestication of *T. turgidum* started with its cultivation in the Fertile Crescent some 10,000 years ago^5^. Subsequent hybridization with *Aegilops tauschii* (DD) formed the hexaploid *T*. *aestivum* (AABBDD)^6^. Whereas ancient gene flow incorporated the majority of the AABB genome diversity into hexaploid wheat, only a small fraction of the D genome diversity was captured^7^. Indeed, hybridization between *T. turgidum* and *Ae. tauschii* was thought to be restricted to a sub-population of *Ae. tauschii* from the shores of the Caspian Sea in present-day Iran^8^. Despite sampling limited diversity, this genomic innovation created a plant more widely adapted to a broad range of environments, and with end-use qualities not found in its progenitors^1^.

The low genetic diversity of the bread wheat D-subgenome has long motivated breeders to recruit diversity from *Ae. tauschii*. The most common route involves hybridization between tetraploid wheat and *Ae. tauschii* followed by chromosome doubling to create synthetic hexaploids^9^. Alternatively, direct hybridization between hexaploid wheat and *Ae. tauschii* is possible. This approach usually requires embryo rescue but has the advantage that it does not disrupt desirable allele combinations in the bread wheat A- and B-subgenomes^10, 11^. Notwithstanding, the products of all these wide crosses require backcrossing to domesticated cultivars to remove unwanted agronomic traits from the wild progenitor and restore optimal end-use qualities. The boost to genetic diversity and resilience therefore comes at a cost to the breeder^9^. However, if haplotypes underlying useful traits could be directly identified in *Ae. tauschii*, this would mitigate a critical limitation in breeding wheat with *Ae. tauschii*; such haplotypes can be tagged with molecular markers for accelerated delivery into domesticated wheat by combining marker-assisted selection^12^ with rapid generation advancement^13^. Furthermore, a gene-level understanding would permit next-generation breeding by gene editing and transformation.

In this study we performed whole-genome shotgun short-read-sequencing on a diverse panel of 242 *Ae. tauschii* accessions. We discovered that an uncharacterized *Ae. tauschii* lineage contributed to the initial gene flow into domesticated wheat, thus broadening our understanding of the evolution of bread wheat. To facilitate the discovery of useful genetic variation from *Ae. tauschii*, we established a *k*-mer-based association mapping pipeline and demonstrated the mobilization of the untapped diversity from *Ae. tauschii* into wheat through the use of synthetic wheats and genetic transformation for biotic stress resistance genes.

## Results

### Multiple hybridization events shaped the bread wheat D-subgenome

We identified a set of 242 non-redundant *Ae. tauschii* accessions with minor residual heterogeneity after short-read sequencing of 306 accessions covering the geographical range spanned by diverse *Ae. tauschii* collections (Fig. 1a; Supplementary Figs. 1-3; Supplementary Tables 1-5). From the accessions sequenced, we generated a *k*-mer matrix representing the genetic diversity in the *Ae. tauschii* species complex, and a single nucleotide polymorphism (SNP) matrix relative to the AL8/78 reference genome^14^ (Supplementary Files 1 and 2).

**Fig. 1.**
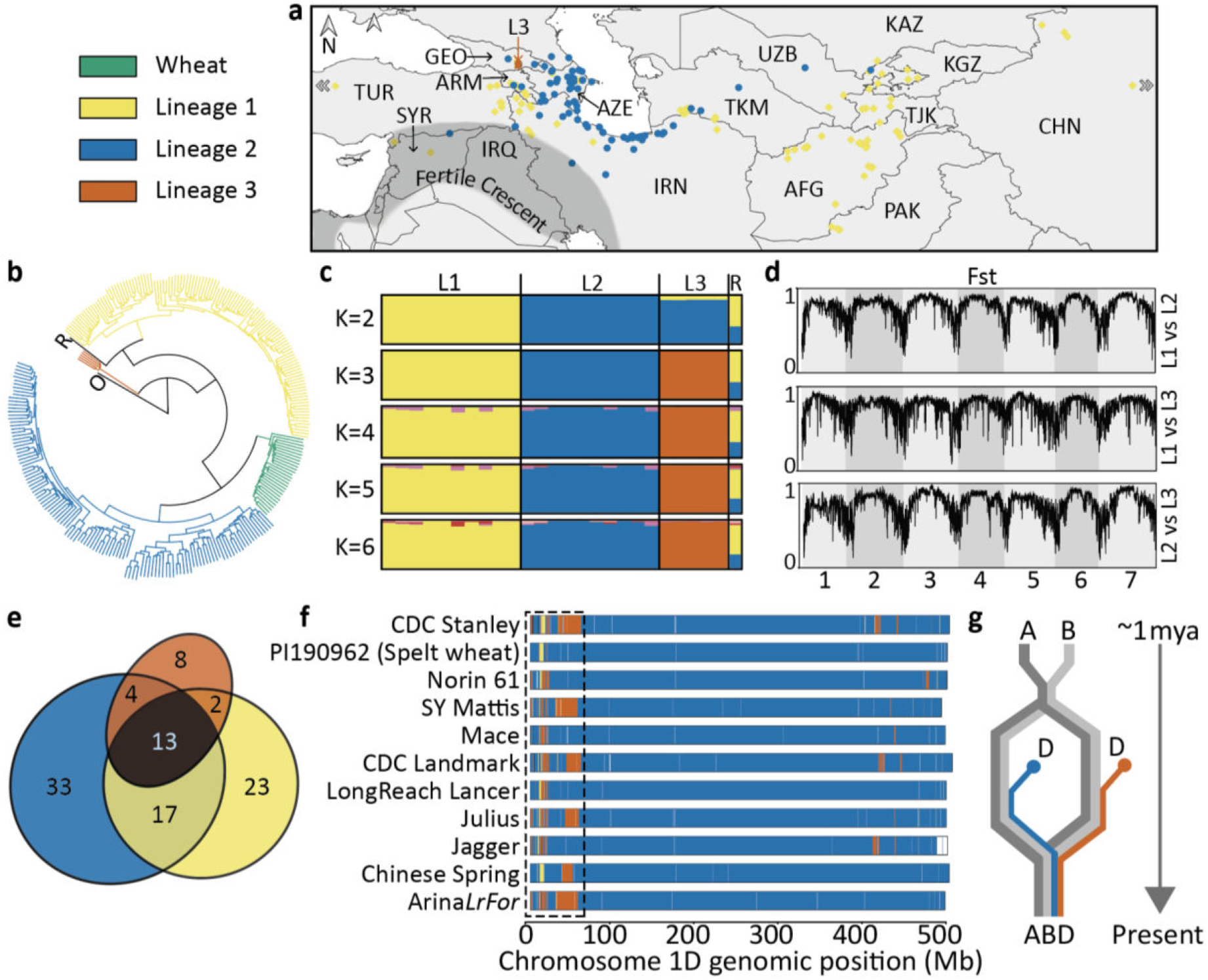
Characterization of a third lineage of *Aegilops tauschii* and its contribution to the wheat D-subgenome. The color code for all panels is shown for wheat and *Ae. tauschii* lineages (L1, L2, L3). **a**, Distribution of 242 *Ae. tauschii* samples used in this study. The five L3 accessions are indicated by an orange vertical arrow. **b**, Phylogeny showing the D-subgenome of 28 wheat landraces in relation to *Ae. tauschii*, a tetraploid (AABB genome) outgroup (O), and an *Ae. tauschii* recombinant inbred line (RIL) derived from Lineages 1 and 2. **c**, STRUCTURE analysis of the randomly selected ten accessions from each of Lineages 1 and 2 along with the five accessions of Lineage 3 and the RIL. **d**, Genome-wide fixation index (F_ST_ ) estimates of the *Ae. tauschii* lineages. **e**, Venn diagram showing percentage of lineage-specific and shared *k*-mers between the lineages. **f**, Chromosome 1D of wheat cultivars/accessions colored according to their *Ae. tauschii* lineage-specific origin. **g**, The pattern of lineage-specific contribution to the wheat D-subgenome (highlighted for one region by a dashed rectangle in panel f) suggests that at least two polyploidization events with distinct *Ae. tauschii* lineages, followed by intraspecific crossing, gave rise to extant hexaploid bread wheat.

*Ae. tauschii* is generally categorized into two lineages, Lineage 1 (L1) and Lineage 2 (L2)^15, 16^, with L2 considered the major contributor to the wheat D-subgenome^8^. To better understand the relationship between *Ae. tauschii* and wheat, we randomly selected 100,000 *k*- mers and checked their presence in the short-read sequences of 28 hexaploid wheat landraces^17^. We used a tetraploid wheat accession as an outgroup in the phylogenetic analysis and included a recent *Ae. tauschii* L1-L2 recombinant inbred line (RIL)^15^ as a control in our population structure analysis. We generated a phylogeny based on the presence/absence of these *k*-mers and found it to be consistent with earlier phylogenies generated using molecular markers (Fig. 1b)^8, 15, 16^. In particular, *Ae. tauschii* L1 and L2 formed two major clades, whereas the wheat D-subgenome formed a discrete and narrow clade most closely related to L2. This supports the L2 origin of the wheat D-subgenome and its limited genetic diversity relative to *Ae. tauschii*. A group of five accessions formed a distinct clade separate from L1 and L2, as previously observed^15, 16^, which seems to be a basal lineage based on the split from the outgroup. Matsuoka *et al.* (2013) hypothesized that this group could be a separate lineage^18^, whereas Singh *et al.* (2019) hypothesized that it could have arisen from inter-lineage hybridization followed by isolated evolution^15^. To resolve this question, we conducted Bayesian clustering analysis using STRUCTURE^19^. Because this algorithm does not reliably recover the correct population structure when sampling is uneven^20^, we randomly selected 10 accessions from each of L1 and L2 for this analysis, along with the five accessions of the putative Lineage 3 (L3) and the control L1-L2 RIL (Supplementary Table 6). Performing STRUCTURE analysis with K=2 showed the putative L3 accessions as an admixture of L1 and L2, similar to the L1-L2 RIL, but with K=3, these accessions were assigned to a distinct lineage (Fig. 1c). Further increasing the value of K did not reveal any discernible sub-structure. This interpretation was supported by the ΔK curve, which showed a clear peak at K=3 (Supplementary Fig. 4). Principle component analysis also separated the population into three clusters corresponding to L1, L2 and L3 (Supplementary Fig. 4). Computing the genome-wide pairwise fixation index (F_ST_ ) between the three lineages using SNPs in a sliding window of 1 Mb with a step size of 100 kb indicated a high level of population differentiation across the genome (Fig. 1d). These observations demonstrate the existence of a differentiated third lineage within *Ae. tauschii*.

Consistent with the above population structure, we found that 64% of the *Ae. tauschii* pan- genome, as represented by *k*-mers, is lineage-specific (Fig. 1e). We used the lineage-specific *k*-mers to understand the origin of the wheat D-subgenome by representing the D-subgenomes of the available chromosome-scale wheat assemblies^21^ as 100 kb segments and assigning them to the *Ae. tauschii* lineage predominantly contributing lineage-specific *k*-mers to that segment (Supplementary Fig. 5). To account for recent alien introgressions in modern cultivars due to breeding, only those *k*-mers which were also present in the 28 hexaploid wheat landraces^17^ were used. The differential presence of L2 and L3 segments at multiple independent regions in these wheat lines (shown for Chromosome 1D in Fig. 1f and Chromosomes 2D-7D in Supplementary Fig. 6) suggests that at least two hybridization events gave rise to the extant wheat D-subgenome (Fig. 1g) and that one of the D genome donors was of predominantly L3 origin. The total L3 contribution across all the seven chromosomes ranges from 0.5% for Spelt to 1.9% for Arina*LrFor*, with an average of 1.1% for all the 11 reference genomes (Supplementary Fig. 6; Supplementary File 3).

### Discovery of *Ae. tauschii* candidate genes underpinning biotic and abiotic resilience, phenology and yield-component traits

Identification of genes or haplotypes in *Ae. tauschii* underpinning useful variation would permit accelerated wheat improvement through wide crossing and marker-assisted selection or biotechnological approaches to introduce them into wheat. To identify this variation, we adapted our *k*-mer-based association mapping pipeline, previously developed for resistance gene families obtained using sequence capture^22^, to whole genome shotgun data (Supplementary Fig. 7a). Instead of mapping the *k*-mers directly to the *Ae. tauschii* AL8/78 reference genome, we generated a *de novo* assembly of accession TOWWC0112 (N50=196 kb), which carries two cloned stem rust resistance genes that could be used as controls, and then anchored this assembly to a reference genome^14^. This enabled identification of the *cis*- associated *k*-mers rather than those linked in repulsion to the corresponding region in the reference genome (Supplementary Fig. 7c-e). We also determined that the sequencing coverage could be reduced from 10-fold to 5-fold with no appreciable loss of signal from the two control genes (Supplementary Fig. 8). To test our method further, we performed association mapping for resistance to additional stem rust isolates and flowering time. For stem rust, we identified a peak within the genetic linkage group of *SrTA1662*^23^ (Fig. 2a; Supplementary Fig. 9; Supplementary Table 7). Annotation of the associated 50 kb linkage disequilibrium (LD) block revealed two genes, of which one encoded the nucleotide-binding and leucine-rich repeat (NLR) gene previously identified in our sequence capture association pipeline^24^ (Fig. 2a; Supplementary Fig. 10; Supplementary Tables 8-9). We also recorded flowering time and found that it mapped to a broad peak of 5.46 Mb on chromosome arm 7DS containing 35 genes including *FLOWERING LOCUS T1* (Fig. 2b; Supplementary Fig. 11; Supplementary Tables 7,9), a well-known regulator of flowering time in dicots and monocots, including wheat^25–27^.

**Fig. 2.**
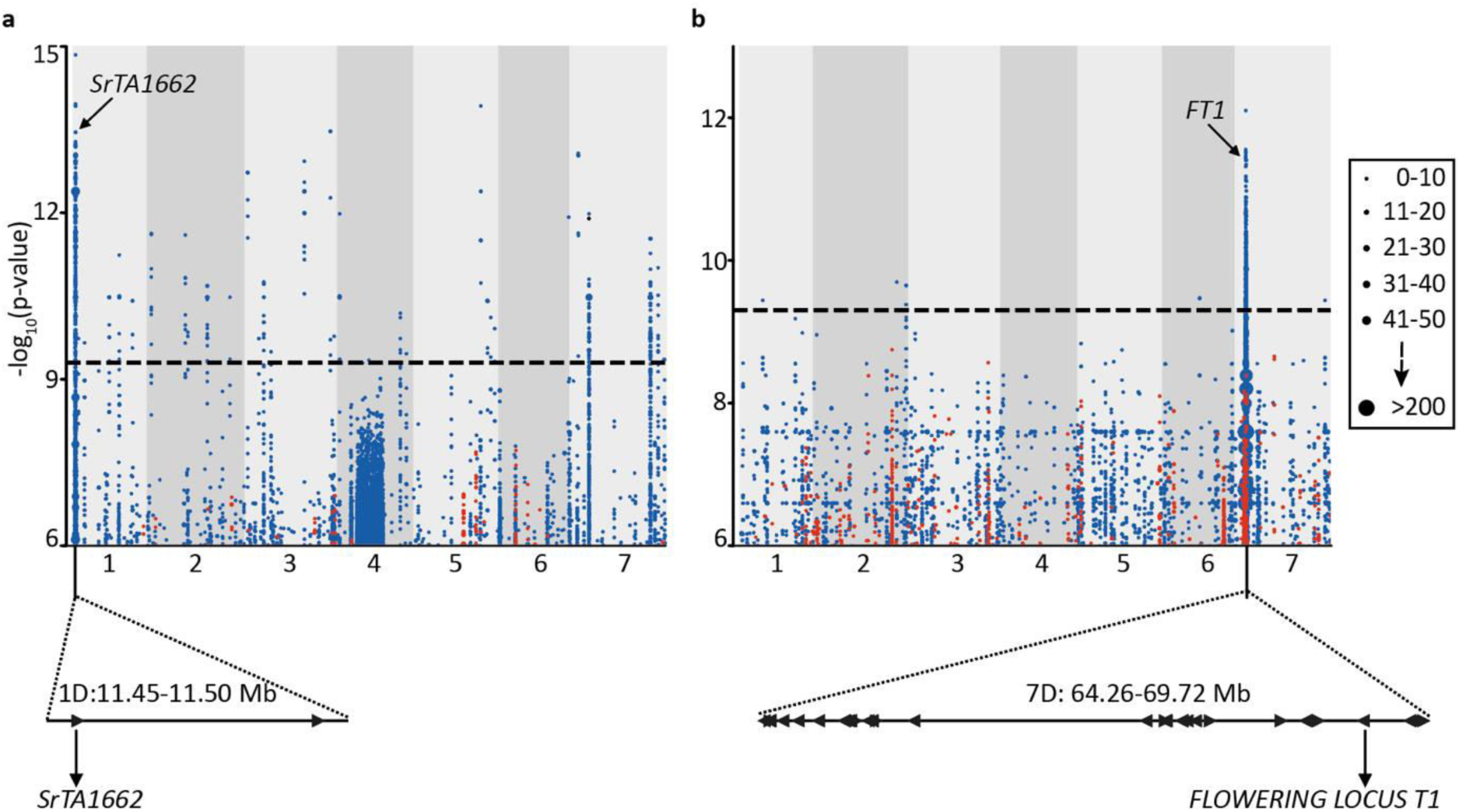
Genetic identification of candidate genes for stem rust resistance and flowering time by *k*-mer–based association mapping. **a**, *k*-mers significantly associated with resistance to *Puccinia graminis* f. sp. *tritici* race QTHJC mapped to scaffolds of a *de novo* assembly of *Aegilops tauschii* accession TOWWC0112 anchored to chromosomes 1 to 7 of the D-subgenome of Chinese Spring^28^. Points on the y-axis show *k*-mers significantly associated with resistance (blue) and susceptibility (red). **b**, *k*-mers significantly associated with flowering time mapped to *Ae. tauschii* reference genome AL8/78 with early (red) or late (blue) flowering time association relative to the population mean across the diversity panel. Candidate genes for both phenotypes are highlighted. Point size is proportional to the number of *k*-mers (see insert).

**Fig. 3.**
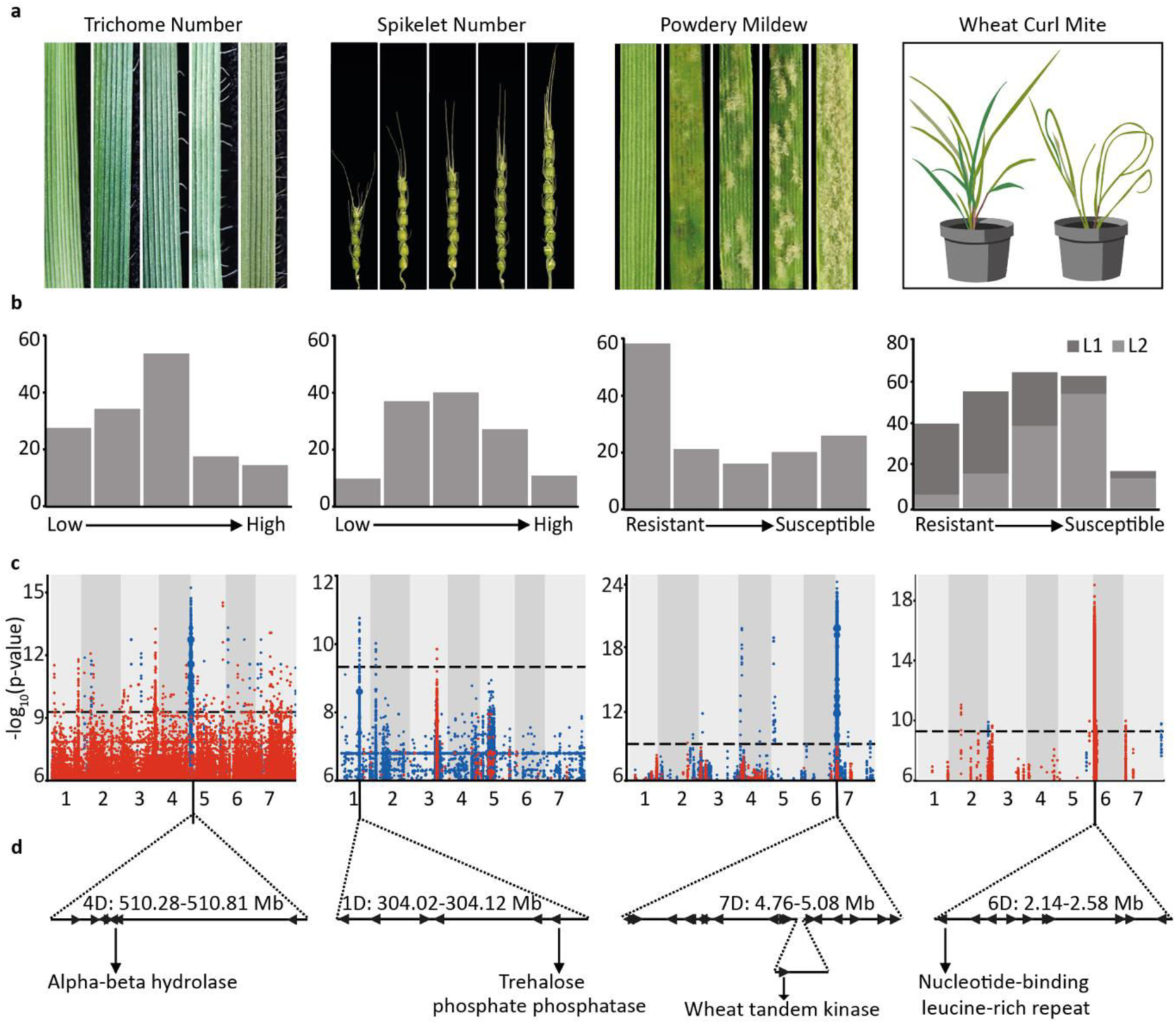
Genome-wide association mapping in *Aegilops tauschii* for morphology, disease and pest resistance traits. **a**, Representation of the scale of phenotypic variation observed. **b**, Frequency distribution of the different phenotypic scales corresponding to panel a. Lineage 1 (L1) and Lineage 2 (L2) are shown in dark and light grey, respectively. **c**, *k*-mer–based association mapping to a *de novo* assembly of accession TOWWC0112 anchored to the AL8/78 reference genome (trichome number, spikelet number) or accession TOWWC0106 anchored to AL8/78 (response to powdery mildew), or directly mapped to AL8/78 (response to wheat curl mite). *k*-mer color-coding and dot size are as in Fig. 2. **d**, Identification of genes under the peak in the GWAS plot with promising candidate(s) indicated. The wheat tandem kinase gene resides within 60 kb insertion relative to the AL8/78 reference genome.

We next screened the *Ae. tauschii* panel for leaf trichomes (a biotic and abiotic resilience trait^29, 30^, spikelet number per spike (a yield component), infection by *Blumeria graminis* f. sp. *tritici* (cause of powdery mildew), and resistance to the wheat curl mite, *Aceria tosichella* (vector of wheat streak mosaic virus)^31^ (Supplementary Table 7). All four phenotypes presented continuous variation in the panel (Fig. 3ab; Supplementary Fig. 12). Mean trichome number along the leaf margin mapped to a 530 kb LD block on chromosome arm 4DL (Fig. 3c; Supplementary Table 9), within a 12.5 cM region previously defined by biparental linkage mapping^32^. The 530 kb interval contains seven genes including an alpha-beta hydrolase, a gene class with significantly increased transcript abundance in developing trichomes of *Arabidopsis thaliana*^33^. The number of spikelets per spike was associated with a discrete 100 kb peak on chromosome arm 1DL containing six genes (Fig. 3b; Supplementary Table 9). One of these encodes a trehalose-6-phosphate phosphatase (TPP) that is homologous to RAMOSA3 and TPP4 known to control inflorescence branch number in maize^34^, and SISTER OF RAMOSA3 that influences spikelet fertility in barley^35^. Powdery mildew resistance mapped to a 320 kb LD block on chromosome arm 7DS containing nineteen genes in the resistant haplotype, including a ∼60 kb insertion with respect to the reference genome AL8/78 (Fig. 3c; Supplementary Table 9). No NLR immune receptor-encoding gene was detected; however, the insertion contains a wheat-tandem kinase (WTK), a gene class previously reported to confer resistance to wheat stripe rust (*Yr15*)^36^, stem rust (*Rpg1* and *Sr60*)^37, 38^ and powdery mildew (*Pm24*)^39^. Resistance to wheat curl mite mapped to a 440 kb LD block on chromosome arm 6DS within a region previously determined by bi- parental mapping^40–42^ (Fig. 3c; Supplementary Table 9). The interval contained ten genes including an NLR immune receptor, a gene class previously reported to confer arthropod resistance in melon and tomato^43^. These results, and a rapid LD decay (Supplementary Fig. 13), highlight the ability of the panel and *k*-mer based association mapping to identify candidate genes, including those in insertions with respect to the reference genome, within discrete genomic regions for quantitative traits of agronomic value.

### *Ae. tauschii* Lineages 1 and 2 share regions of low genetic divergence

We investigated the population-wide distribution of the candidate genes controlling disease resistance and morphology identified by association mapping (Figs. 2-3) across a genome-wide phylogeny of *Ae. tauschii* and a worldwide collection of 28 wheat landraces^17^. The absence of the alleles promoting disease resistance, more spikelets and higher trichome density in the wheat landraces for the novel candidate genes suggest they were not incorporated into the initial gene flow into wheat (Fig. 4a). We next examined the distribution of these alleles between the three lineages of *Ae. tauschii*. The *Cmc4* gene candidate for resistance to wheat curl mite was largely confined to L1, whereas the allele variants promoting higher trichome density, spikelet number and resistance to wheat stem rust and powdery mildew were largely confined to L2 (Fig. 4a). Exceptions included three occurrences of the *Sr46* gene in L1 and five occurrences of the candidate *Cmc4* gene in L2. To investigate whether this was due to a common genetic origin or convergent evolution, we generated phylogenies based on the SNPs within the respective 200 kb and 440 kb *Sr46* and *Cmc4* LD blocks. This showed that all functional haplotypes clustered together irrespective of genome-wide lineage assortment, indicative of a common genetic origin, and not convergent evolution (Fig. 4bc; Supplementary Table 10).

**Fig. 4.**
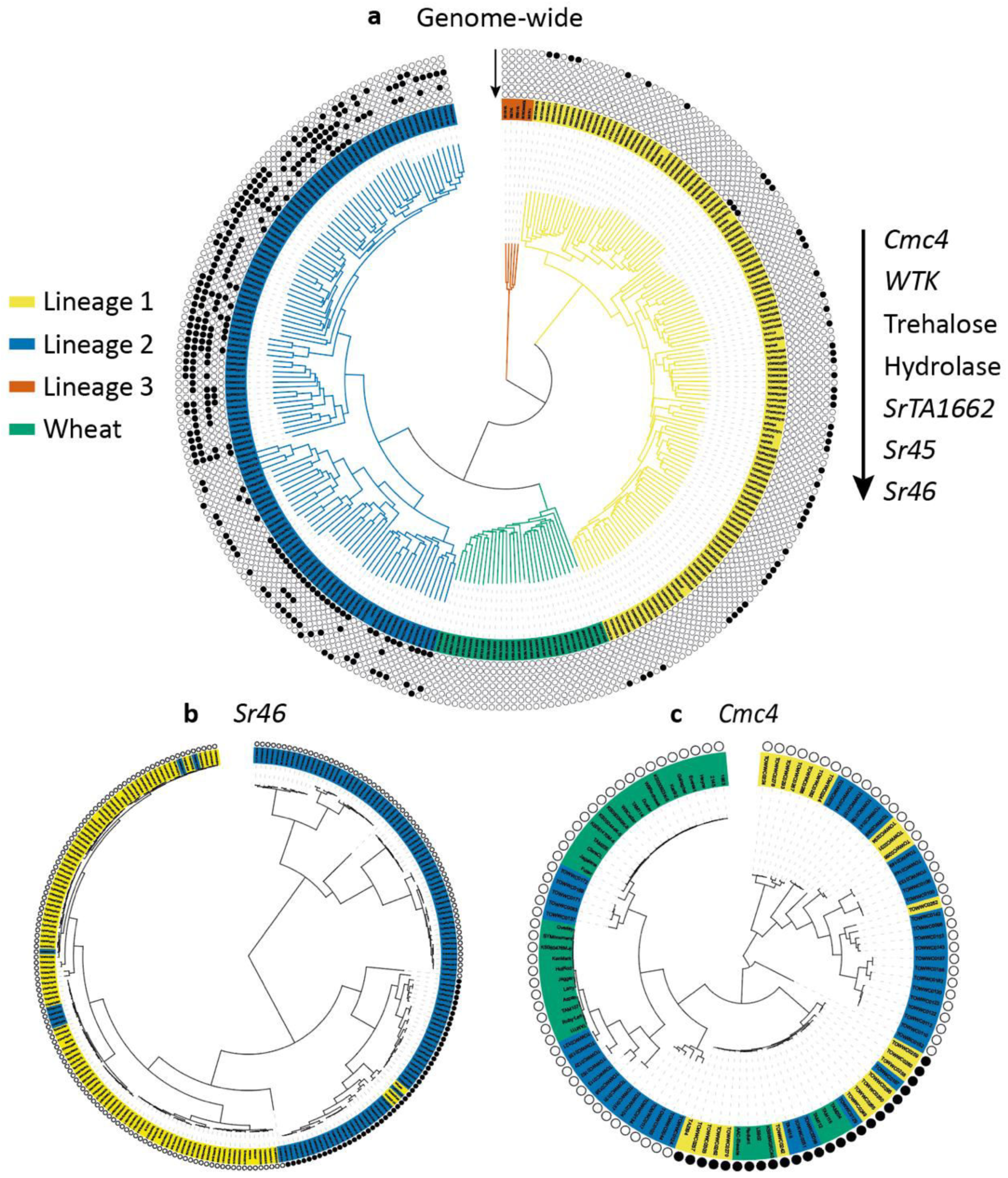
Genome-wide and gene-level relationship between *Aegilops tauschii* and wheat. **a**, Genome-wide *k*-mer-based phylogeny of *Ae. tauschii* and hexaploid wheat landraces with designation of presence of candidate and cloned genes/alleles for disease and pest resistance, and morphological traits based on a genome-wide phylogeny. Presence and absence of allele-specific polymorphisms is indicated by circles filled with black or white, respectively. **b**, Phylogeny of *Ae. tauschii* L1 and L2 accessions based on SNPs restricted to the 200 kb region surrounding *Sr46*. **c**, Phylogeny based on SNPs of the 440 kb region in LD with *Cmc4*. Only the most resistant and susceptible *Ae. tauschii* accessions were included, along with resistant and susceptible modern elite wheat cultivars (different from the landraces shown in panel a).

### Post-domestication mobilization of *Ae. tauschii* genes into wheat

The ability to precisely identify *Ae. tauschii* haplotypes and candidate genes for target traits provides an opportunity for accelerating their introduction into cultivated wheat. We selected 32 non-redundant and genetically diverse *Ae. tauschii* accessions, which capture 70% of the genetic diversity across all lineages, and crossed them to tetraploid durum wheat (*Triticum turgidum* var. *durum*; AABB) to generate independent synthetic hexaploid wheat lines (Fig. 5. Supplementary Table 11). From this ‘library’, we selected four synthetic lines with the powdery mildew *WTK* candidate resistance gene. These synthetics as well as their respective *Ae. tauschii* donors were resistant to powdery mildew, while the durum line was susceptible (Fig. 6a; Supplementary Fig. 14). Annotation of *WTK* identified seven alternative transcripts of which only one, accounting for ∼80% of the transcripts, leads to a complete 2,160 bp 12- exon open reading frame (Fig. 6a; Supplementary Fig. 15; Supplementary Tables 12-13). Next, we targeted two exons with very low homology to other genes for virus-induced gene silencing (VIGS). *WTK*-containing *Ae. tauschii* and synthetics inoculated with the *WTK*- VIGS constructs became susceptible to powdery mildew whereas empty vector-inoculated plants remained resistant (Fig. 6a; Supplementary Fig. 14). This supports the conclusion that *WTK,* hereafter designated *WTK4,* is required for powdery mildew resistance and remains effective in synthetic hexaploids. Thus, these synthetic lines can serve as pre-breeding stocks for introduction of the trait into elite wheat.

**Fig. 5.**
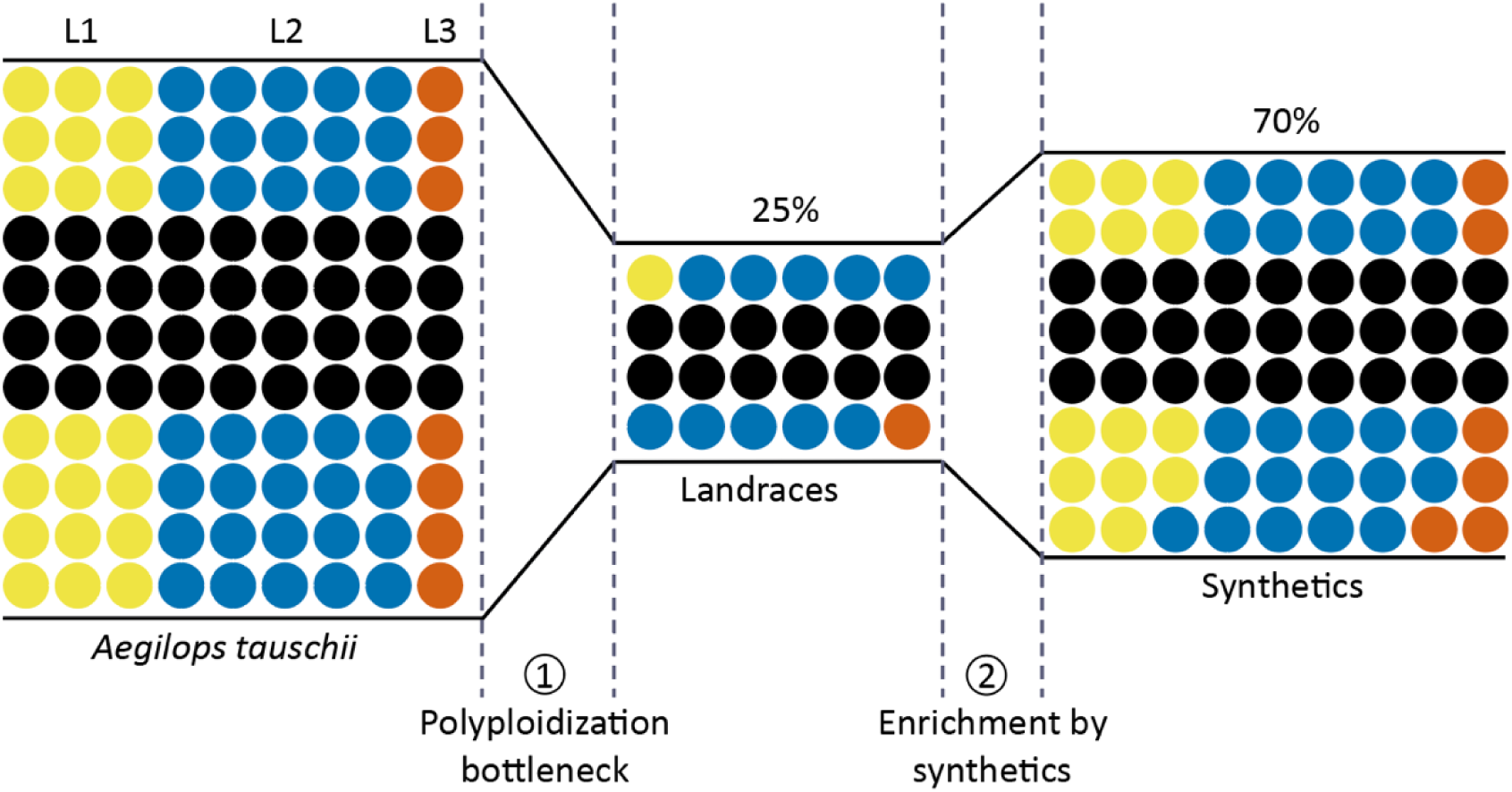
Restricted gene flow from *Aegeilops tauschii* to wheat and the capture of *Ae. tauschii* diversity in a panel of synthetic hexaploid wheats. Genetic diversity private to *Ae. tauschii* Lineages 1, 2 and 3 is color-coded blue, red and green, respectively, whereas black dots represent *k*-mer sequences (51-mers) common to more than one lineage. The number of dots is proportional to the number of *k*-mers. The polyploidization bottleneck (1) incorporated 25% of the variant *k*-mers found in *Ae. tauschii* into wheat landraces. The addition of 32 synthetic hexaploid wheats (2) restored this to 70%.

**Fig. 6.**
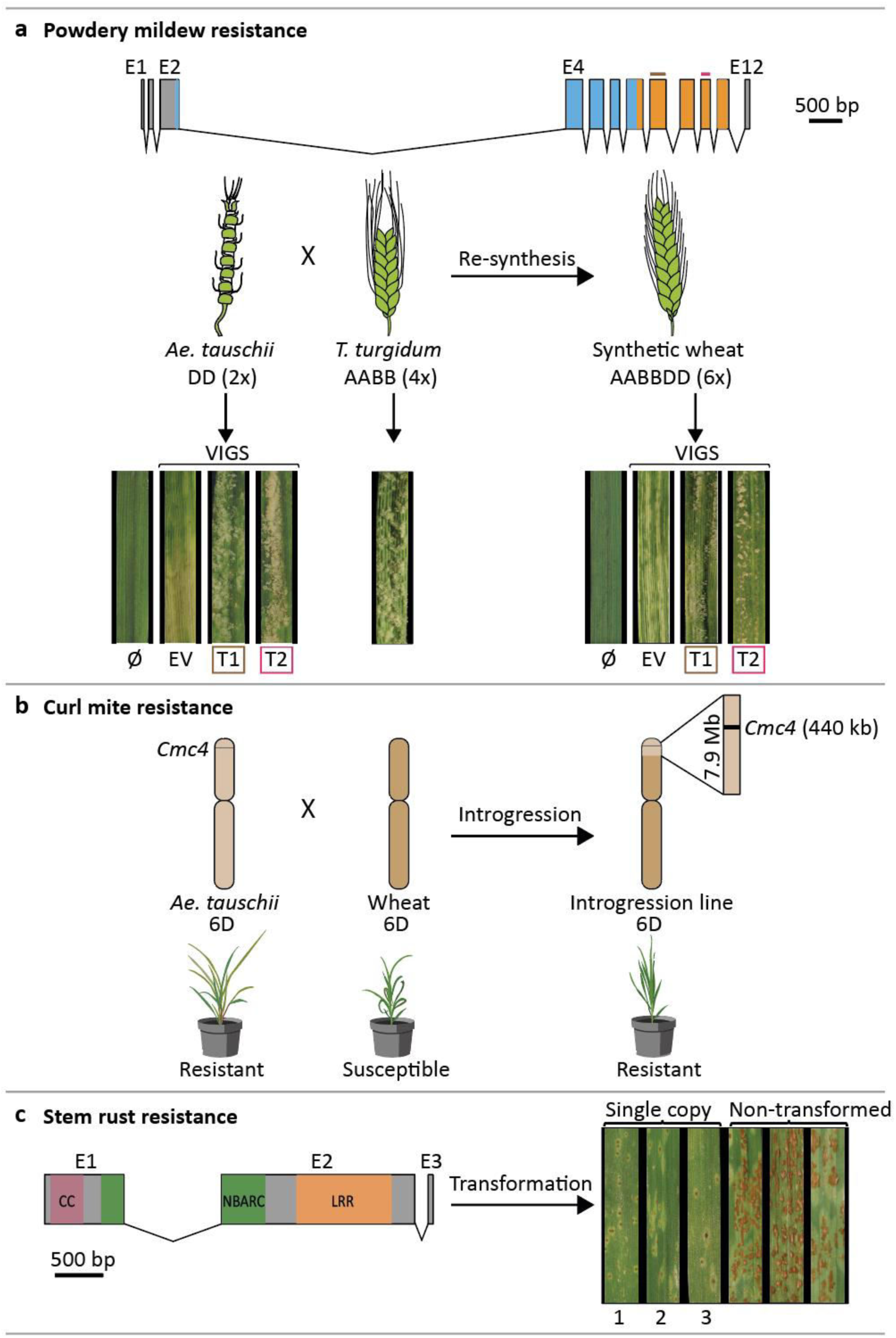
Functional transfer of disease and pest resistance from *Aegilops tauschii* into wheat. **a**, *WTK4* gene structure represented by rectangles (exons, E1 to E12) joined by lines (introns). Kinase domains are shown in blue and orange. Exons used for designing virus- induced gene silencing (VIGS) target 1 (T1) and target 2 (T2) are shown in brown and red, respectively. Below, schematic of the cross between *Ae. tauschii* accession Ent-079 (contains *WTK4*) and *Triticum turgidum* durum line Hoh-501 (lacks *WTK4*) that generated the synthetic hexaploid wheat line NIAB-144. Leaf segments from plants subjected to VIGS with empty vector (EV), T1, T2, or non-virus control (Ø) and super-infected with *Blumeria graminis* f. sp. *tritici* isolate *Bgt96224* avirulent to *WTK4*. **b**, Introgression of the *Cmc4* locus from *Ae. tauschii* accession TA1618 into wheat. The 440 kb *Cmc4*linkage disequilibrium block (black) resides within a 7.9 Mb introgressed segment on chromosome 6D (light brown) in wheat cultivar ‘TAM 115’. Below, drawings of wheat curl mite-induced phenotypes. **c**, Structure of the *SrTA1662* candidate gene. The predicted 970 amino acid protein has domains with homology to a coiled-coil (CC), nucleotide-binding (NB-ARC) and leucine-rich repeats (LRR). Right, transformation with an *SrTA1662-*genomic construct into cv. Fielder and response to *Puccinia graminis* f. sp. *tritici* isolate UK-01 (avirulent to *SrTA1662*) of single- copy hemizygous transformants (1, DPRM0059; 2 DPRM0051; 3, DPRM0071) and non- transgenic controls.

Developing wheat cultivars improved with traits from *Ae. tauschii* can also be achieved by direct crossing between the diploid and hexaploid species^10^. The wheat curl mite resistance genes *Cmc4* was originally transferred by crossing of *Ae. tauschii* accession TA2397 (L1) into wheat^44, 45^, and genetically localized to chromosome 6D in agreement with our association mapping^40, 41,42^. Given the common resistant haplotype of *Cmc4* in L1 and L2 (Fig. 4), we hypothesized that *Cmc4* is the same as a gene originating from L2 accession TA1618, which was introgressed at the same locus into wheat cv. ‘TAM 112’ via a synthetic wheat^41, 45^. Consistent with this hypothesis, we observed the same haplotype at the wheat curl mite resistance locus across all derived resistant hexaploid wheat lines and in the *Ae. tauschii* donors of *Cmc4* and *Cmc_TAM112_* (Fig. 4). We delimited the length of the introgressed *Ae. tauschii* wheat curl mite fragments by comparing SNP data for resistant wheat lines and the corresponding *Ae. tauschii* donors. The TA2397 (L1) introgression spanned 41.5 Mb whereas the TA1618 (L2) introgression was reduced to 7.9 Mb in wheat cv. ‘TAM 115’ (Fig. 6b; Supplementary Fig. 16).

As an alternative to conventional breeding, we targeted the *SrTA1662* candidate stem rust resistance gene (Fig. 2d) for introduction into wheat by direct transformation. We cloned a 10,541 bp genomic fragment encompassing the complete *SrTA1662* transcribed region, as well as >3 kb of 3’ and 5’ UTR putative regulatory sequences; this was sufficient to confer full race-specific stem rust resistance in transgenic wheat (Fig. 6c; Supplementary Fig. 17; Supplementary Table 14).

## Discussion

The origin of hexaploid bread wheat has long been the subject of intense scrutiny. Archaeological and genetic evidence suggests that diploid and tetraploid wheats were first cultivated 10,000 years ago in the Fertile Crescent (Fig. 1a)^5, 6^. The expansion of tetraploid wheat cultivation northeast into Caspian Iran and towards the Caucasus region resulted in sympatry with *Ae. tauschii* and the emergence of hexaploid bread wheat^6^. *Ae tauschii* displays a high level of genetic differentiation among local populations and genetic marker analysis suggests that the wheat D-subgenome donor was recruited from an L2 population of *Ae. tauschii* in the southwestern coastal area of the Caspian Sea^8^. However, not all the diversity within the wheat D-subgenome can be explained by a single hybridization event^6, 46, 47^. Our population genomic analysis revealed the existence of a third lineage of *Ae. tauschii*, L3, which also contributed to the extant wheat genome. For example, a glutenin allele required for superior dough quality was recently found to be of L3 origin^48^. L3 accessions are restricted to present-day Georgia and may represent a relict population from a glacial refugium as observed in Arabidopsis^49^. We observed genomic signatures specific to L2 and L3 in hexaploid wheat supporting the multiple hybridization hypothesis (Fig 1g).

The creation of hexaploid bread wheat, whilst giving rise to a crop better adapted to a wider range of environments and end uses^1^, came at the cost of a pronounced genetic bottleneck^7^. Our analysis suggested that only 25% of the genetic diversity of *Ae. tauschii* contributed to the initial geneflow into hexaploid wheat (Fig. 5). To explore this diversity, we performed association mapping and discovered new gene candidates for disease and pest resistance and agro-morphological traits underpinning abiotic stress tolerance and yield, exemplifying the potential of *Ae. tauschii* for wheat improvement (Fig. 6). We obtained discrete LD blocks of 50 to 520 kb, with the exception of flowering time, which resulted in a broad LD block of 5.5 Mb around the *FT1* locus (Fig. 2). The low degree of historical recombination around *FT1* is likely imposed by the reduced probability of intra-species hybridization between populations carrying alleles promoting different flowering times. In contrast to the discrete mostly sub-megabase mapping intervals we obtained by association mapping with *k*-mer-based marker saturation, conventional biparental mapping studies on the D-subgenome resulted in large intervals with a median of 10 Mb (Supplementary Table 15).

In polyploid wheat recessive variants are not readily observed, hence genetics and genomics in wheat has mostly focused on rare dominant or semi-dominant variants^50^. Reflecting this, out of 64 genes cloned in polyploid wheat by forward genetics, at least 58 have dominant or semi-dominant modes of action (Supplementary Table 16). This constraint is removed in *Ae. tauschii* by virtue of being diploid, which along with its rapid LD decay, makes it an ideal platform for gene discovery by association mapping. Genes and allelic variants discovered in *Ae. tauschii* can subsequently be studied in wheat by generating transgenics or mutants. Moreover, our public library of synthetic wheats, which captures 70% of the diversity present across all three *Ae. tauschii* lineages, allows immediate trait assessment in a hexaploid background. The trait-associated haplotypes can be used to design molecular markers to precisely track the desired gene in a breeding program. In conclusion, our study provides an end-to-end pipeline for rapid and systematic exploration of the *Ae. tauschii* gene pool for improving modern bread wheat.

## Methods

### Compilation of an inventory of *Ae. tauschii* accessions

We surveyed the literature, germplasm banks and unpublished collections to compile an inventory of 641 *Ae. tauschii* potentially unique accessions spanning its geographical range from western Turkey to eastern China. Accordingly, accessions of *Ae. tauschii* described by Singh *et al.* (2019) (549 accessions)^51^, Arora *et al.*, (2019) (150 accessions)^22^ and Abbasov *et al.* (2019) (36 accessions)^52^ were cross-referenced to identify duplicates (accessions with the same name, or identical accessions with different names). Where available, missing collection site data from these lines was obtained from www.genesys-pgr.org. Within this set, genotyping-by-sequencing data^51^ or records held at the Wheat Genetics Resource Centre, Kansas State University, USA, were used as further criteria to highlight genetically redundant lines. To this set, we added the geographic data from an unpublished collection of 81 accessions established by Ali Mehrabi at the Ilam University Gene Bank (IUGB). This collection includes 63 accessions collected across the moderate climatic ranges of the Zagros and Alborz mountains in northern and western Iran in 2006. Each sample was isolated based on a separation of at least 20 km or by geographical barriers. The spike samples from an area within a radius of 500 m were collected as one population (accession). Spikelet shape was used to identify subspecies, with intermediate forms considered as ssp. *tauschii*. Samples were bulked at Ilam University Research Farm, deposited to the Cereal Gene Bank at Ilam University, Īlām, Iran and assigned unique IUGB germplasm identification codes. An additional collection of 84 L1 accessions from Tajikistan, collected between 2005 and 2006 by Firuza Nasyrova, Institute of Botany, Plant Physiology and Genetics, Tajik National Academy of Sciences Tajikistan, was also included^53^. These accessions were assigned latitude and longitude coordinates based on the 13 geographical locations listed. In total, the inventory includes 641 accessions, of which passport data is available for 588 accessions (Supplementary Table 1; Supplementary Fig. 1). Passport data for the sequenced D donors was also compiled as above, with further latitude and longitude data to that described in Arora *et al.* (2019) provided by Huw Jones, NIAB or obtained from www.genesys-pgr.org (Supplementary table A; Supplementary Fig. I).

### Selection of accessions for whole genome shotgun short-read sequencing

Previously available KASP genotyping data from Arora *et al.* (2019)^22^ was used to select a set of 24 genetically diverse *Ae. tauschii* accessions for short read Illumina sequencing with an average coverage of 30. The criteria used for selecting accessions included representation of different branches of a phylogenetic tree^22^, country of origin, disease variation for six races of stem rust^54^, and accessions carrying mapped and/or cloned resistance genes from *Ae. tauschii* (*Sr45*, *Sr33*, *Sr46*, *SrTA1662*, *Lr21* and *SrTA10171*). Of these 24 accessions, seven were from L1, sixteen from L2, and one from L3. Of the 171 accessions remaining in the diversity panel of the Arora *et al.* (2019) study^22^, 130 L2 accessions were selected to maximize non-redundancy based on the histogram of similarity scores obtained from the KASP analysis (Supplementary Fig. 2). A threshold of 97.5% separated the peak of high similarity values and was therefore taken to be the redundancy cut-off. Notwithstanding this cut-off, among the 130 selected accessions, 28 accessions were above the redundancy cut-off but were included due to secondary considerations including disease variation for stem rust races and seed availability. These 130 accessions were targeted for whole genome shotgun (WGS) sequencing with an average coverage of 10.

To cover the genetic diversity of L1, we selected 110 putative L1 accessions based on the analysis performed by Singh *et al.* (2019)^51^. These accessions were also targeted for WGS sequencing at 10-fold genome coverage. Two of these accessions (TOWWC0253, line BW_23897 and TOWWC0304, line BW_23925) were subsequently found to belong to L2 after analysis of the sequencing data.

After discovering the distinct signature of L3 in wheat, 10 putative inter-lineage hybrids from Singh *et al*. (2019)^51^ were sequenced with an average coverage of 7.5-fold genome coverage. Seven of these putative hybrids were found to belong to L3, one was found to belong to L1, and, as previously determined by Singh et al. (2019)^51^, two were recent (possibly, post collection) recombinant inbred lines of L1 and L2.

We also sequenced 40 *Ae. tauschii* donors used to create synthetic hexaploid wheats with the durum wheat cultivar Hoh-501 (see below). Eleven of these donors were included in the above selections. The remaining 29 were sequenced with an average coverage of 7.5-fold.

In addition, we sequenced accessions TA1618 and TA2394, which were used for introgression of *Cmc4* into wheat, giving a tally of 306 sequenced accessions which were used for subsequent analyses. Separately, we included two more L2 accessions TA2450 and Clae 23, the former because of the strategic importance as the parent of a TILLING population^55^. Based on the GBS and KASP analyses, respectively, referred to above, these accessions are redundant in the panel (Supplementary Table 1).

### DNA extraction and WGS-sequencing of the *Ae. tauschii* diversity panel and the durum wheat cultivar Hoh-501

DNA from accessions TOWWC0001 to TOWWC0195 was made available from Arora *et al.* (2019)^22^. For accessions TOWWC0196 to TOWW0304, the SHW lines and their diploid and tetraploid donors, high molecular weight DNA was extracted following a modified CTAB protocol^56^. For accessions TOWW0196 to TOWW0304 the same plant was used for DNA and single seed descent. Accessions TOWWC0001 to TOWW0304, and the SHW diploid and tetraploid donors, were sequenced on an Illumina platform with 150 bp PE chemistry at Novogene (China). The targeted sequencing coverage ranged from 7-fold (39 accessions), to 10-fold (234 accessions) to 30-fold (24 accessions) (Supplementary Table 2).

### RNA extraction and RNAseq

Total RNA was obtained from accessions TOWWC0020 (line BW_20689), TOWWC0104 (line BW_21074), TOWWC0106 (line BW_21086), TOWWC0107 (line BW_21090), TOWWC0108 (line BW_21096), TOWWC0112 (line BW_21114), TOWWC0134 (line BW_21215) and TA2450. Seeds were germinated in a Petri dish at 22 °C in the dark in the presence of 0.5 ppm gibberellic acid and then transferred to a cereal soil mix^13^ and placed in a controlled environment room at 18 °C with a 16 hour light/ 8 hour dark cycle. Leaf tissue from two-week old seedlings was harvested in the afternoon and immediately used for RNA extraction using a Qiagen RNA extraction kit followed by bead size selection. The RNA was used to generate Illumina TruSeq libraries which were sequenced on an Illumina platform with 150 bp paired end reads to generate the following amounts of data per accession: TOWWC0020, 34.98 Gb; TOWWC0104, 12.58 Gb; TOWWC0106, 34.47 Gb; TOWWC0107, 12.46 Gb; TOWWC0108, 35.36 Gb; TOWWC0112, 38.28 Gb; TOWWC0134, 33 Gb; TA2450, 35 Gb.

### SNP-calling relative to the AL8/78 reference genome

Following WGS sequencing, we called SNPs across the panel relative to the *Ae. tauschii* AL8/78 reference genome assembly. The 306 *Ae. tauschii* samples were aligned to the *Ae. tauschii* AL8/78 reference genome^14^ using hisat2 default parameters. All alignment bam files were sorted and duplicates removed using samtools (view -bhs and otherwise default). All bam files were fed into the variant call pipeline using bcftools (-q 20 -a DP,DV | call -mv -f GQ) with parallelization ‘-r $region’ of 4 Mb windows for a total of 1010 intervals (regions). The raw variant files were filtered or recalled using a published AWK script based on DP/DV ratios with default parameters (https://bitbucket.org/ipk_dg_public/vcf_filtering/src/master/) except minPresent parameter (we used minPresent=0.8 and minPresent=0.1). The minPresent=0.8 dataset was used for redundancy analysis. The minPresent=0.1 and minPresent=0.8 were both used for GWAS analysis. The resulting matrix (104 million SNPs for minPresent=0.1) were uploaded to Zenodo.

### Quality control for genetic redundancy and residual heterogeneity of sequenced *Ae. tauschii* accessions

A total of 100,900 (100 every 4 Mb window) SNPs were randomly chosen to compute pairwise identity by state (IBS) among all samples for a total of 46,665 comparisons using custom R and AWK scripts (https://github.com/wheatgenetics/owwc). For every sample pair, a percent identity greater than 99.5% was deemed redundant based on the histogram distribution of all IBS values (Supplementary Fig. 2). This analysis confirmed the results of the KASP analysis conducted on the L2 accessions above (Supplementary Fig. 2).

For each accession (except TOWWC0193, which is related to the reference genome AL8/78), the fraction of heterozygous SNPs in the total number of bi-allelic SNPs was computed. Based on the distribution of these values (Supplementary Fig. 3; Supplementary Table 3), 0.1 was deemed to indicate a low degree of residual heterogeneity. BW_26042, with a value of 0.17, was found to be the only outlier exceeding this threshold.

Based on these QC analyses, a non-redundant and genetically stable set of 242 accessions was retained for further analysis. The redundant pairs, along with the different similarity scores, are given in Supplementary Table 4 and the set of 242 non-redundant accessions is provided in Supplementary Table 5.

### *De novo* assembly of non-redundant accessions from whole genome shotgun short-read data

The primary sequence data of non-redundant accessions were trimmed using Trimmomatic v0.238 and *de novo* assembled with the MEGAHIT assembler using default parameters^57^. The output of the assembler for each accession was a FASTA file containing all the contig sequences. The assemblies are available from Zenodo.

### Genome sequencing and assembly of *Ae. tauschii* accession TOWWC0112 (line BW_01111)

TOWWC0112 was assembled by combining paired-end and mate-pair sequencing reads using TRITEX^58^, an open-source computational workflow. A PCR-free 250 bp paired-end library with an insert size range of 400-500 bp was sequenced to a coverage of ∼70. Mate-pair libraries MP3 and MP6, with insert size ranges of 2-4 kb and 5-7 kb, respectively, were sequenced to a coverage of ∼20. The assembly generated had an N50 of 196 kb. The assembly is available from the electronic Data Archive Library (e!DAL).

### Genome sequencing and assembly of *Ae. tauschii* accession TOWWC0106 (line BW_01105)

Accession TOWWC0106 was sequenced on the PacBio Sequel II platform (Pacific Biosciences) with single molecule, real-time (SMRT) chemistry and on the Illumina platform. For SMRT library preparation, ∼7 μg of high-quality genomic DNA was fragmented to 20 kb target size and assessed on an Agilent 2100 Bioanalyzer^59^. The sheared DNA was end repaired, ligated to blunt-end adaptors, and size selected. The libraries were sequenced by Berry Genomics (Beijing, China). A standard Illumina protocol was followed to make libraries for PCR-free paired-end genome sequencing with ∼1 μg of genomic DNA that was fragmented and size-selected (350 bp) by agarose gel electrophoresis. The size-selected DNA fragments were end-blunted, provided with an A-base overhang, and then ligated to sequencing adapters. A total of 251.8 Gb high-quality 150 paired-end (PE150) PCR-free reads were generated and sequenced on the NovaSeq sequencing platform.

A set of 11.35 million PacBio long reads (289.6 Gb), representing a ∼66-fold genome coverage was assembled using the CANU pipeline with default parameters^60^. The assembled contigs were polished with 251.8 Gb PCR free reads using Pilon default parameters^61^. The resulting assembly had an N50 of 1.5 Mb. The assembly is available from e!DAL.

### Annotation of the TOWWC0112 and TOWWC0106 genome assemblies

Structural gene annotations for TOWWC0112 (line BW_01111) and TOWWC0106 (line BW_01105) were done combining two annotation strategies: prediction based on comparative *ab initio* gene finding and a lift-over approach. The comparative *ab initio*approach warrants identification of gene model variations, while ensuring uniformity between the annotated genomes. To counteract any further intrinsic error during the *ab initio* gene calling, a consolidation step by projecting the existing *Ae. tauschii* gene models onto the BW_01111 and BW_01105 scaffolds was applied.

Comparative *ab initio* gene prediction resorts to utilising whole-genome sequence alignments (WGA). So, a WGA between TOWWC0112, TOWWC0106 and *Ae. tauschii*^14^ using the cactus pipeline (Version 1.0)^62^ was constructed. Prior to the alignment step, all nucleotide sequences were 20-kmer-softmasked to reduce complexity and facilitate construction of the WGA using the tallymer subtools from the genome tools package (Version 1.6.1)^63^. The resulting WGA was used as an input for the Augustus comparative annotation pipeline (Version 3.3.3)^64^. Extrinsic hints for intron and CDS regions were generated from the *Ae. tauschii* annotation in order to guide gene calling and reduce false positive predictions. Furthermore, a previously trained specific Aegilops model was used.

Predicted gene models were subsequently classified into high- or low-confidence. Non-redundant candidate protein sequences were compared against the following three manually curated databases using BLASTp: first, PTREP, a database of hypothetical proteins that contains deduced amino acid sequences in which, in many cases, frameshifts have been removed, which is useful for the identification of divergent TEs having no significant similarity at the DNA level; second, UniPoa, a database comprised of annotated Poaceae proteins; third, UniMag, a database of validated magnoliophyta proteins. UniPoa and UniMag protein sequences were downloaded from Uniprot and further filtered for complete sequences with start and stop codons. Best hits were selected for each predicted protein to each of the three databases. Only hits with an E-value below 10e-10 were considered. A high confidence protein sequence is complete and has a subject and query coverage above the set threshold of 66% in the UniMag database, or no blast hit in UniMag but in UniPoa and not TREP. A low confidence protein sequence is not complete and has a hit in the UniMag or UniPoa database but not in TREP, or no hit in UniMag and UniPoa and TREP but the protein sequence is complete.

For a consistent annotation between BW_01111, BW_01105 and *Ae. tauschii* AL8/78, gene models from *Ae. tauschii* AL8/78 were additionally projected onto the BW_01111 and BW_01105 sequences using a lift-over approach described in^65^. Afterwards, high-confidence *ab initio* and projected gene models were combined. In case of overlap, the *ab initio* gene model was retained. The resulting merger was again confidence-classified using the methods described above.

On top of this homology-based classification, further tweaking was done relying on functional assignment. First, functional annotations of predicted protein sequences were generated using the AHRD pipeline (https://github.com/groupschoof/AHRD). Second, human-readable description lines were scanned for TE, plastid and non-protein coding keywords and gene models were tagged accordingly. Any non-tagged low-confidence proteins with an AHRD 3*-rating were promoted to high-confidence. Contrarily, high-confidence proteins were demoted if their AHRD rating was only a one star. An overview of the number of high confidence and low confidence annotated genes can be found in Supplementary Table 8.

Completeness of the predicted gene space was measured with BUSCO (version 4.06, viridiplantae orthodb10)^66^ (Supplementary Fig. 10). The annotations are available from e!DAL.

Phenotyping the *Ae. tauschii* diversity panel and SHW lines

### Wheat stem rust

The wheat stem rust phenotypes with *P. graminis* f. sp. *tritici* isolate 04KEN156/04, race TTKSK, and isolate 75ND717C, race QTHJC, were obtained from the study of Arora *et al.* (2019)^22^. As part of this study, we also phenotyped the same *Ae. tauschii* lines, with isolate UK-01 (race TKTTF)^67^ (Supplementary Table 7) using the same procedures as described in^68^.

### Trichomes

For counting trichomes and measuring flowering time in *Ae. tauschii*, 50 L1 accessions and 150 L2 accessions were pre-germinated at ∼ 4°C in Petri dishes on wet filter paper for two days in the dark. They were transferred to room temperature (∼20°C) and daylight for four days. Three seedlings of each genotype were transplanted on January 22, 2019 into 96-cell trays filled with a mixture of peat and sand, and then grown under natural vernalization in a glasshouse with no additional light source or heating at the John Innes Centre, Norwich, UK. Trichome phenotyping was conducted one month later. Close-up photographs of the second leaf from seedlings at the three-leaf stage were taken, visualized in ImageJ, and the number of trichomes were counted along one side of a 20 mm leaf margin in the mid-leaf region. Measurements were taken from three biological replicates (Supplementary Table 7).

### Flowering time, biological replicate 1

Three seedlings used for trichome phenotyping (see above) were transferred on 25 March into individual 2 litre pots filled with cereal mix soil^69^. Flowering time was recorded when the first five spikes were three-quarters emerged from the flag leaf sheath, equivalent to a 55 on the Zadoks growth scale^70^ (Supplementary Table 7).

### Flowering time, biological replicates 2 and 3

A total of 147 *Ae. tauschii* L2 accessions were grown in the winters of 2018/19 and 2019/20 in the greenhouse at the Department of Agrobiotechnology, University of Natural Resources and Life Sciences, Vienna, Austria. Seeds of each accession were sown in multi-trays in a mixture of heat-sterilized compost and sand and stratified for one week before germination at 4°C with a 12 hour day/night light regime. Thereafter, the seeds were germinated at 22°C and at the one leaf stage vernalized for 11 weeks. Five seedlings per accession were transplanted to 4 litre pots (18 cm diameter, 21 cm height) filled with a mixture of heat-sterilized compost, peat, sand and rock flour. In the winter of 2018/19, one pot (= one replicate) per accession was planted, whereas in 2019/20, two pots (= two replicates) were planted. The pots were randomly arranged in the greenhouse and maintained at a temperature of 14/10°C day/night with 12 hour photoperiod for the first 40 days. At spike emergence, the temperature was increased to 22/18°C day/night with 16 h photoperiod at 15,000 lx. At least ten spikes per pot were evaluated for beginning of anthesis, taken as 60 on the Zadoks growth scale^70^, resulting in a minimum of 30 assessed spikes per accession. Flowering time was recorded every second day.

The flowering date was analyzed using a linear mixed model, which considered subsampling of individual spikes within each pot as follows:

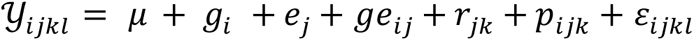

where 𝒴_*ijkl*_ denotes the flowering date observation of the individual spikes, *μ* is the grand mean and *g*_*i*_ is the genetic effect of the *i*th accession. The environment effect *e*_*j*_ is defined as the effect of the *j*th year and the genotype by environment interaction is described by *ge*_*ij*_. *r*_*jk*_ is the effect of the *k*th replication within the *j*th year, *p*_*ijk*_ the effect of the ith pot within the *k*th replication and *j*th year, and *ε*_*ijkl*_ is the residual term. Analysis was performed with R 3.5.1 (R Core team 2020)^71^ using the package *sommer*^72^ with all effects considered as random, except *g*_*i*_ which was modelled as a fixed effect to obtain the best linear unbiased estimates (BLUEs) (Supplementary Table 7).

### Spikelets per spike

For *Ae. tauschii* spikelet phenotyping, 151 accessions from L2 were vernalized at constant 4°C for eight weeks in a growth chamber (Conviron, Winipeg, Canada). After vernalization, the accessions were transplanted to one-gallon pots in potting mix (peat moss and vermiculite) and placed in a temperature-controlled Conviron growth chamber with diurnal temperatures gradually changing from 12°C at 02:00 to 17°C at 14:00 with a 16 h photoperiod and 80% relative humidity. To represent biological replication each accession was grown in two pots, and each pot contained two plants. At the transplanting stage, 10 g of a slow-release N-P-K fertilizer was added to each pot. At physiological maturity, 5-15 main stem/tiller spikes per replication (*i.e.* per pot) were harvested and the number of immature, as well as mature, spikelets counted. Any obvious weak heads from late-growing tillers were not included. Least square means for each replication were used for *k*-mer-based association genetic analysis (Supplementary Table 7).

### Powdery mildew

Resistance to *Blumeria graminis* f. sp. *tritici* was assessed with *Bgt96224*, a highly avirulent isolate from Switzerland^73^, using inoculation procedures previously described^74^. Disease levels were assessed 7-9 days after inoculation as one of five classes of host reactions: resistance (R; 0-10% of leaf area covered), intermediate resistance (IR; 10- 25% of leaf area covered), intermediate (I; 25-50% of leaf area covered), intermediate susceptible (IS; 50-75 % of leaf area covered) and susceptible (S; >75% of leaf area covered) (Supplementary Table 7).

### Wheat curl mite

A total of 210 *Ae. tauschii* accessions, 102 from L1 and 108 form L2 (Supplementary Table 7), were screened for their response against wheat curl mite. *Aceria tosichella* (Keifer) biotype 1 colonies (courtesy of Dr Michael Smith, Department of Entomology, Kansas State University) were mass-reared under controlled conditions at 24°C in a 14 h light, 10 h dark cycle, using the susceptible wheat cv. Jagger. The biotype 1 colony was previously reported as avirulent towards all *Cmc* resistance genes^40, 75–77^. A single colony consisted of an individual pot with ∼50 seedlings and 20 colonies were grown to have sufficient mite inoculum to conduct the phenotyping. Colonies were placed inside 45 cm x 45 cm x 75 cm mite-proof cages covered with a 36 µm mesh screen (ELKO Filtering Co., Zurich, Switzerland) to avoid contamination until being used to infest the *Ae. tauschii* accessions. Accessions from L1 and L2 were evaluated in independent experiments. Six plants per accession were individually grown in 5 cm x 5 cm x 5 cm pots under controlled conditions at 24°C in a 14 h day, 10 h dark cycle. Pots were arranged randomly in an incomplete block design where the block was the tray fitting 32 pots (8 rows and 4 columns). A single pot with the susceptible check cv. Jagger was included in each tray. Accessions were infested at the two-leaf stage with mite colonies collected from infested pieces of leaves from the susceptible plants and spread as straw over the pots. Plants were evaluated individually 10-14 days after infestation. Wheat curl mite damage was assessed as curled or trapped leaves using a visual scale from 0 to 4, with 0 no symptoms and 1 to 4 increasing levels of curliness or trapped leaves (Supplementary Fig. 12).

The adjusted mean or best linear unbiased estimator (BLUE) for each accession was calculated with the ‘lme4’ R package^78^ using a linear regression model as:

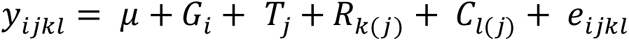

where *y*_*ijkl*_ is the phenotypic value, µ is the overall mean, *G*_*i*_ is the fixed effect of the *i*^*th*^ accession (genotype), *T*_*j*_ is the random effect of the *j*^*th*^ tray assumed distributed as iid 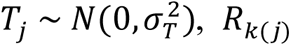 is the random effect of the *k*^*th*^ row nested within the *j*^*th*^ tray assumed distributed as iid 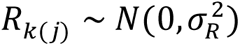, *Rk*(*j*) is the random effect of the *k*^th^ column nested within the *j*^*th*^ tray assumed distributed as iid 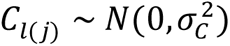, and *e*_*ijkl*_ is the residual error distributed as iid 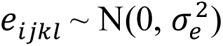.

### *k*-mer presence/absence matrix

*k*-mers (k = 51) were counted in trimmed raw data per accession using Jellyfish^79^. *k*-mers with a count of less than two in an accession were discarded immediately. *k*-mer counts from all accessions were integrated to create a presence/absence matrix with one row per *k*-mer and one column per accession. The entries were reduced to 1 (presence) and 0 (absence). *k*- mers occurring in less than two accessions or in all but one accession were removed during the construction of the matrix. Programs to process the data were implemented in Python and are published at https://github.com/wheatgenetics/owwc. The k-mer matrix is available from e!DAL.

### Phylogenetic tree construction

A random set of 100,000 *k*-mers was extracted from the OWWC *k*-mer matrix to build a UPGMA (unweighted pair group method with arithmetic mean) tree with 100 bootstraps using the Bio.Phylo module from the Biopython (http://biopython.org) package. Further, a Python script was used to generate an iTOL (https://itol.embl.de/) compatible tree for rendering and annotation. The Python script and the random set of 100,000 *k*-mers used for generating the tree is available at https://github.com/wheatgenetics/owwc.

### Bayesian cluster analysis using STRUCTURE

Bayesian clustering implemented in STRUCTURE^19^ version 2.3.4 was used to investigate the number of distinct lineages of *Ae. tauschii*. To control the bias due to the highly unbalanced proportion of the three groups^20^ in the non-redundant sequenced accessions (119 accessions of L2, 118 accessions of L1 and five accessions of putative L3), 10 accessions each of L1 and L2 were randomly selected for each STRUCTURE run, along with the five accessions of the putative L3 and the control L1-L2 RIL. The random selection of 10 accessions each of L1 and L2 was performed 11 times without replacement, thus covering a total of 110 accessions each of L1 and L2 over 11 STRUCTURE runs (Supplementary Table 6). STRUCTURE simulations were run using a random set of 100,000 *k*-mers with a burn-in length of 100,000 iterations followed by 150,000 Markov Chain Monte-Carlo iterations for five replicates each of K ranging from 1 to 6. STRUCTURE output was uploaded to Structure Harvester (http://taylor0.biology.ucla.edu/structureHarvester)80 to generate a ΔK plot for each run. For each STRUCTURE run, a clear peak was observed at K=3 in the ΔK plot, suggesting that there are three distinct lineages of *Ae. tauschii*^19, 80^. STRUCTURE results were processed and plotted using CLUMPAK^81, 82^ to maintain the label collinearity for multiple replicates of each K.

### Determination of genome-wide fixation index

Genome-wide pairwise fixation index (F_ST_ ) between the three *Ae. tauschii* lineages was computed using VCFtools^83^ with parameters “--fst-window-size” and “--fst-window-step” set to 1,000,000 and 100,000, respectively.

### Assignment of wheat D-subgenome segments to *Ae. tauschii* lineages

For each of the 11 chromosome-scale wheat assemblies^21^, *k*-mers were considered as usable only when present at a single locus in the D-subgenome. Furthermore, out of these *k*-mers, for nine modern cultivars, only those *k*-mers were considered usable which were also present in the short-read sequences from 28 hexaploid wheat landraces^17^. For the assembled wheat genomes, each chromosome of the D-subgenome was divided into 100 kb non-overlapping segments. A 100 kb segment was assigned to *Ae. tauschii* if at least 20% of 100,000 *k*-mers within that segment were usable as well as present in at least one non-redundant *Ae. tauschii* accession. A segment assigned to *Ae. tauschii* was further assigned to one of the three lineages – L1, L2, L3 – if the count of usable *k*-mers specific to that lineage exceeded the count of those specific to the other lineages by at least 0.01% of 100,000 *k*-mers. Scripts to determine the counts of lineage-specific and total *Ae. tauschii k*-mers per 100 kb segment are published at https://github.com/wheatgenetics/owwc and the output files obtained for 11 wheat assemblies are collated in an Excel file which is available from Zenodo.

### Anchoring of a *de novo* assembly to a reference genome

The contigs of a *de novo* assembly were ordered along a chromosome-level reference genome using the minimap2 alignment tool^84^, and assigning them the genomic coordinates of their longest hits.

### Correlation pre-filtering

For each of the assembly *k*-mers, if also present in the pre-calculated presence/absence matrix, Pearson’s correlation between the vector of that *k*-mer’s presence/absence and the vector of the phenotype scores was calculated. Only those *k*-mers for which the absolute value of correlation obtained was higher than a threshold (0.2, by default) were retained in order to reduce the computational burden of association mapping using linear regression.

### Linear regression model accounting for population structure

To each filtered *k*-mer from the previous step, a *p*-value was assigned using linear regression with a number of significant PCA dimensions as covariates to control for the population structure. PCA was computed using the aforementioned set of 100,000 *k*-mers. The exact number of significant PCA dimensions was chosen heuristically. Too high a number might over-correct for population structure, while too few might under-correct. In the context of this study, three dimensions were found to represent a good trade-off.

### Approximate Bonferroni threshold computation

For each phenotype in this study, the total number of *k*-mers used in association mapping varied between 3,000,000,000 and 5,000,000,000. In general, if the *k*-mer size is 51, a SNP or any other structural variant would give rise to at least 51 *k*-mer variants. Therefore, the total number of tested *k*-mer variants should be divided by 51 to get the effective number of variants to adjust the *p*-value threshold for multiple testing. Assuming a *p*-value threshold of 0.05, a Bonferroni-adjusted negative log *p*-value threshold between 9.1 and 9.3 was obtained for each phenotype. The more stringent cut-off of 9.3 was chosen throughout this study.

### Generating association mapping plots

Association mapping plots were generated using Python. For a chromosome-level reference assembly, each integer on the x-axis corresponds to a 10 kb genomic block starting from that position. For an anchored assembly, each integer on the x-axis represents the scaffold which is anchored starting from that position. Dots on the plot represent the negative log *p*-values of the filtered *k*-mers within each block. Dot size is proportional to the number of *k*-mers with the specific negative log *p*-value. The plotting script is published at https://github.com/wheatgenetics/owwc.

Optimizing *k*-mer GWAS in *Ae. tauschii* with the positive controls *Sr45* and *Sr46*

We used previously generated stem rust phenotype data for *Puccinia graminis*f. sp. *tritici* isolate 04KEN156/04, race TTKSK, on 142 *Ae. tauschii* L2 accessions^24^. Mapping *k*-mers with an association score >6 to the *Ae. tauschii* reference genome AL8/78 gave rise to significant peaks for the positive controls *Sr45* and *Sr46* (Supplementary Fig. 7a). The peaks contain *k*-mers which are negatively correlated with resistance (shown as red dots) because the AL8/78 reference accession does not contain *Sr45* and *Sr46*. To identify the true *Sr45* and *Sr46* haplotypes, accession TOWWC0112 (which contains *Sr45* and *Sr46*)^22^ was assembled from 10-fold whole genome shotgun data using MEGAHIT (N50 1.1 kb) and used in association mapping. However, noise masked the positive signals from *Sr45* and *Sr46* when the short scaffolds were distributed randomly along the x-axis (Supplementary Fig. 7b). Anchoring the scaffolds to the AL8/78 reference genome considerably improved the plot and produced positive signals for *Sr45* and *Sr46* (blue peaks; Supplementary Fig. 7c). An improved assembly (N50 196 kb) generated with mate pair libraries and again anchored to AL8/78, further reduced the background noise (Supplementary Fig. 7d).

### Performing *k*-mer GWAS in *Ae. tauschii* with reduced coverage datasets

The trimmed sequence data of each non-redundant accession was randomly subsampled to reduce the coverage to 7.5-fold, 5-fold, 3-fold and 1-fold. For each coverage point, the *k*-mer GWAS pipeline was applied and *k*-mers with an association score >6 were mapped to the *Ae. tauschii* reference genome AL8/78 (Supplementary Fig. 8).

### Alignment of whole-genome sequence data and data filtering, variant calling, genome-wide association study based on SNP data, and delineation of *Cmc4* region

We performed alignment (bowtie2 v2.2.9 –end-to-end –very-fast) of WGS data to reference AL8/78 and performed variant calling using BCFtools (v1.9) as described above in the genetic redundancy section. We then conducted SNP-based association mapping with Q+K model using GAPIT (10.1093/bioinformatics/bts444) for the wheat curl mite trait. The region was preliminarily determined to be 2.1-2.6 Mb (chr6D) based on a p-value cut-off of 3.7e-7 (FDR = 0.05). We then constructed a more diverse panel including hexaploid wheat (with known wheat curl mite response) to further refine the region and infer allele introgressions. WGS sequence data for 121 entries, 36 wheat lines and a subset of 85 *Ae. tauschii* accessions, 25 from L1 (14 resistant and 11 susceptible) and 60 from L2 (six resistant and 54 susceptible), were aligned to an *in silico* synthetic reference genome created by combining the A and B genomes from the hexaploid wheat cv. Jagger^21^ and the *Ae. tauschii* genome assembly (Aet v4.0; NCBI BioProject PRJNA341983) as the D genome. The *Ae. tauschii* accessions were selected to include all the accessions with conclusive resistant (WCM infestation < 0.07) and susceptible phenotypes (WCM infestation > 3) (Supplementary Table 10). The alignment steps (hisat2) and variant calls are the same as described for the genetic redundancy section. To obtain high-quality SNP markers, VCF files were filtered by read depth (DP>=4) and quality (QUAL >30) using bcftools v1.9^85^, and for minor allele frequency (MAF>=0.1), missing data (minPresent >15%), and heterozygosity (het percent < 5%) using a customized pipeline. Retained SNP markers were 947,937 for chromosome 6DS (0 – 230 Mb) and 3,122 for the WCM resistance interval (1.9 – 2.7 Mb, with an extra 0.1 – to 1.2 Mb flanking based on the region identified above).

### Computing genome-wide linkage disequilibrium

The *Ae. tauschii* AL8/78 reference genome was partitioned into five segments (R1, R2a, C, R2b, R3; Supplementary Fig. 13) based on the distribution of the recombination rate, where the boundaries between these regions were imputed using the boundaries established for the Chinese Spring RefSeqv1.0 D subgenome^28^. PopLDdecay^86^ with the parameter “-MaxDist” set to 5 Mb was used to determine the LD decay in these regions for both L1 and L2. For L2, the value of mean r^2^ in the telomeric regions, R1 and R3, dropped below 0.1 at genomic distances of 291 kb and 476 kb, respectively, while for L1, the corresponding genomic distances were 661 kb and 561 kb, respectively.

### Delimitation of the introgressed segment carrying curl mite resistance in hexaploid wheat

To delimit the extent of the *Ae. tauschii* introgression conferring resistance to wheat curl mite into hexaploid wheat we manually explored SNP polymorphisms using whole genome shotgun data for four wheat curl mite resistant wheat lines, ‘KS96WGRC40’, ‘TAM 112’, ‘TAM 115’, and ‘TAM 204’, and two *Ae. tauschii* resistant donor accessions, TA2397 and TA1618 (Supplementary Figure 16). The variant calling (bcftools) and filtering steps are the same as described above. The ‘KS96WGRC40’ resistance donor is the L1 accession TA2397 and the original line where *Cmc4* was mapped^40, 44^. The ‘TAM 112’ resistance donor is the L2 accession TA1618 (PI-268210) through the cultivar ‘Largo’, and the line where *Cmc_TAM112_* was mapped^41, 45^. ‘TAM 115’ and ‘TAM 204’^87^ are both resistant through ‘TAM 112’. Of particular note is that both wheat lines ‘KS96WGRC40’ and ‘TAM 112’ also carry the *Ae. tauschii* L2 accession TA2460 (BW_01148), susceptible to WCM.

### Gene structure and alternative splicing of *WTK4*

A first *in silico* annotation of the *WTK4* gene was done based on RNAseq data from the *Ae. tauschii* accession TOWWC0107 (line BW_01106). We used the SMARTer^TM^ RACE cDNA Amplification Kit (634923, Clontech) to determine the 5’- and 3’–ends using *WTK4*-specific primers (Supplementary Table 12) on cDNA generated from TOWWC0112 mRNA, which was extracted with the Dynabeads™ mRNA DIRECT™ Purification Kit (61012, Invitrogen). For reverse transcription of cDNA, the 3’ SMART CDS Primer II A was replaced by primer GH438 in the 5’ RT reaction. Subsequently, the same reaction containing the tailed first strand cDNA could be used for both 3’ and 5’ race PCR. 5’ RACE PCR was performed with 2 µl of 1:5 diluted cDNA in a 20 µl reaction with the KAPA2G Robust PCR Kit (KK5501, Sigma-Aldrich, St. Louis, Missouri, USA) buffer B, gene specific reverse primer JS633 and the UPM primer provided with the Kit. 30 cycles were run according to the touchdown PCR program 1 described in the SMARTer™ RACE Kit manual. The 3’ race PCR was made with 4 µl of 1:5 diluted cDNA in a 20 µl reaction with the KAPA2G Robust PCR Kit and buffer B, gene specific forward primer JS655 and the universal reverse primer GH439. After initial denaturation at 95°C for 3 min, a touchdown PCR protocol with 10 cycles of 95°C for 15 s, 68°C (-0.8°C/cycle) for 30 s, 72°C for 30 s, then 25 cycles at 95°C for 15 s, 61°C for 15 s, 72° for 30 s was performed with a final extension at 72° C for 5 min. 3’ and 5’ race PCR fragments were subjected to agarose gel electrophoresis, excised, cloned and then sequenced by Sanger sequencing to determine the UTRs. Based on 5’ RACE reactions, we could confirm the presence of at least 82 bp of 5’ UTR without alternative start codons and a 3’ UTR of at least 155 bp.

Guided by the 5’ and 3’ UTRs, we designed primers located on both UTRs to study gene structure and splicing. *WTK4* transcript accumulation was confirmed by PCR amplification using the primers JS693xJS671 followed by a semi-nested reaction with primers JS696xJS671 using KAPA Hifi HotStart Polymerase (KK2502, Kapa Biosystems, Hoffmann-La Roche) with an annealing temperature of 60°C and extension time of 3 min. PCR products were subcloned using CloneJET PCR Cloning Kit (K1232, Thermo Fischer Scientific, Waltham, Massachusetts, USA), according to the manufacturer’s recommendations, and transformed into *E. coli*. Fifty-one single-colony derived plasmid clones were sequenced with the internal primers JS655, JS656, JS661, JS678, JS682, JS689, JS833 and JS835 (Supplementary Table 12). Transcript 1, which encodes the complete WTK4 protein, was found to be the most common transcript (80%), while the other six isoforms are less abundant (between 2% and 10%) and encode truncated proteins (Supplementary Fig. 15; Supplementary Table 13). Core kinase domains were predicted based on the Conserved Domain Database (CDD) from NCBI^88^.

### Virus-induced gene silencing of *WTK4*

To minimize the possibility of off-target silencing, we blasted the *WTK4* coding sequence against the reference genome assemblies of wheat cv. Chinese Spring^28^ and *Ae. tauschii*^14^ for the selection of *WTK4* gene fragments of 150-250 bp with no homology to other genes. Primers JS657xJS658 and JS662x663, with *Not*I and *Pac*I restriction sites in antisense direction to enable antisense insertion in the pBS-BSMV-γ vector, were used to amplify WTK4_target_1 (Exon 10) and WTK4_target_2 (Exon 8), respectively. Seeds from selected *Ae. tauschii* and synthetic hexaploid wheat accessions were stratified at 4°C for five days and then placed in a growth chamber (Conviron, Winnipeg, Canada) cycled at 23°C/16°C, 16/8 hour photoperiod with 60% humidity and a light intensity regime of 350 μmol/s·m^2^. An equimolar amount of pBS-BSMV-α, pBS-BSMV-β and pBS-BSMV-γ transcripts carrying WTK4_target_1 or WTK4_target_2, was inoculated into fully-expanded first leaves of selected *Ae. tauschii* and synthetic hexaploid wheat accessions, and the durum donor Hoh- 501, as previously described^89–91^. For *in vitro* synthesis of viral RNA, the Invitrogen™ mMESSAGE mMACHINE™ T7 Transcription Kit (Thermo Fischer Scientific, Waltham, Massachusetts, USA) was used according to the manufacturer’s recommendations. 14 days after virus infection, 3^rd^ and 4^th^ leaves were detached, placed on 0.5% agar plates complemented with 10 g/L Benzylaminopurine and infected with *Bgt96224*. Powdery mildew phenotypes were visually assessed 7-10 days after infection (Supplementary Fig. 14).

### Gene structure of *SrTA1662* and annotation of its predicted protein

The *SrTA1662* gene was assembled using overlapping contigs from assemblies of *Ae. tauschii* accessions TOWWC0107 (line BW_01106) and TOWWC0050 (line BW_01049) to 10,445 bp. This sequence was confirmed by Sanger-sequencing of overlapping PCR fragments generated from genomic DNA. The intron-exon structure of the gene was predicted by mapping the RNAseq data from TOWWC0107 (line BW_01106). The gene was annotated to have three exons that encode a 970 amino acid protein with a CC and NB-ARC domain and 14 LRRs predicted by NCBI and Pfam databases and LRRpredictor^92^ (Fig. 6c).

### Engineering of *SrTA1662* binary construct for transformation

A 10,455 bp fragment encompassing 3,395 bp of putative 5’ regulatory sequence, 4,026 bp from the ATG to the STOP, and 3,0724 bp of putative 3’ regulatory sequence, and flanked by synthetic 5’ *Not*I and 3’ *Pme*I sites, respectively, was synthesized by Thermofisher Scientific. The DNA fragment was cloned into pMA to yield the toolkit vector 18ACBRCC_SRTA1662_pMA, available from Addgene (<u>#154372</u>). The GoldenGate (MoClo) compatible level 2 vector pGoldenGreenGate-M (pGGG-M)^93^ was modified to receive this fragment. Initially, a vector was assembled with the hygromycin phosphotransferase (*hpt*) gene with the castor bean catalase (*CAT-1*) intron driven by the rice *Actin1* promoter and an adaptor at MoClo position 2 which contained the unique restriction enzyme sites *Not*I and *Pme*I. In brief, the Level 1 construct pICH47802- RActpro::HptInt::NosT (selectable maker) and the Postion 2 *Not*I/*Pme*I adaptor were cloned into the binary level 2 vector pGGG-M using standard Golden Gate MoClo assembly^94^. The resulting vector was deemed pGGG-AH-*Not*I/*Pme*I. The native promoter, coding region and 3’ UTR of *SrTA1662* was cloned as one 10,461 bp fragment into the *Not*I and *Pme*I sites of pGGG-AH-*Not*I/*Pme*I using standard cloning procedures, resulting in the final wheat transformation vector pGGG-AH-SrTA1662. The construct was electroporated into the hypervirulent *Agrobacterium tumefaciens* (strain AGL1)^95^ in addition to the helper plasmid pAL155 which contains an additional *Vir*G gene. *Agrobacterium*-standard inoculums^96^ were prepared as previously described^93^.

### Wheat transformation

The hexaploid wheat cv. Fielder, which is susceptible to *P. graminis* f. sp. tritici isolate UK- 01^97^, was transformed as previously described^93^. Under aseptic conditions wheat immature embryos were isolated, pre-treated by centrifugation, inoculated with *Agrobacterium* AGL1 containing pGGG-AH-SrTA1662 and co-cultivated for 3 days. Wheat callus induction and proliferation, shoot regeneration and rooting were carried out under a stringent hygromycin selection regime before the regenerated plantlets were transferred from *in vitro* to soil and acclimatized to ambient conditions. Transgenesis and transgene copy number analysis was performed by iDNA Genetics, Norwich, UK using Taqman qPCR and the *hpt* probebar^93, 98^. Homozygous single-copy T_2_ lines and their respective nulls are available from the Germplasm Resources Unit www.SeedStor.ac.uk under entry numbers DPRM0050 to DPRM0074 (Supplementary Table 14).

### Phenotyping of *SrTA1662* wheat transgenics for stem rust resistance

Primary (T_0_) hemizygous transgenic plants and non-transgenic controls, which had also undergone tissue culture, were phenotyped with *Puccinia graminis* f. sp. tritici isolate UK-01 as previously described^68^, and the infection phenotypes were correlated with the presence of the *hpt* transgene (Fig. 6c; Supplementary Table 14).

To test for race-specificity, three independent T_2_ homozygous lines and their respective non-transgenic segregants (nulls) were phenotyped with four phylogenetically distinct isolates of stem rust from Clade I (isolate KE184a/18, also known as Ug99, race TTKTT), Clade IV-B (isolate ET11a/18, race TKTTF), Clade IV-F (isolate, IT200a/18, race TKKTF), and Clade III-B (isolate IT16a/18, race TTRTF) according to nomenclature implemented by the Global Rust Reference Center (GRRC), Denmark (Supplementary Fig. 17; Supplementary Table 14). The macroscopic phenotype of the lines was investigated on the 1^st^ and 2^nd^ leaf at the seedling stage in a quarantine greenhouse at the GRRC. Four seedlings of each line were grown in peat moss at 20°C for 11 days in two replicates. Urediniospores of the isolates were retrieved from -80°C storage, heat-shocked at 43°C for 5 minutes, and then suspended in light industrial mineral oil (Novec 7200) prior to spray inoculation of plants on day 11. Inoculated plants were incubated in a humid chamber at 18±2°C in darkness overnight and then maintained in a glasshouse at 20±2°C with 16 hour light/ 8 hour darkness. Qualitative infection types (IT) for individual lines and isolates were assessed 15 days post inoculation using a standard 0-4 scale^99, 100^. Infection types up to 3- were considered incompatible (‘resistant’ host), while 3+ and 4 represented a compatible interaction (‘susceptible’ host).

### Creation of synthetic hexaploid wheat lines

A collection of 429 *Ae. tauschii* accessions representing 14 geographic regions was sourced from seven international germplasm collections. Each accession was genotyped using 15 D genome specific microsatellite simple sequence repeat (SSR) markers covering the entire D genome except the 7D chromosome^101^. A subset of 232 accessions were further genotyped at 62 D genome specific SNP loci identified by the UK wheat SNP consortium^102, 103^. A targeted set of 100 individuals were selected which captured both the geographic distribution and genetic diversity of the wider collection to use as D genome donors in a synthetic resynthesis programme. Hybridization compatibility with the tetraploid donors meant we were able to produce fertile hexaploid synthetic lines from 52 of the selected *Ae. tauschii* accessions. Forty-three of these lines were made with the *T. turgidum* var. durum line Hoh-501 (a winter durum line obtained from Friedrich Longin, University of Hohenheim, Germany) and have been described as part of this study.

Seeds of the selected *Ae. tauschii* accessions and the durum line Hoh-501 were sown directly into Levington’s M2 compost and germinated in a controlled glasshouse with a 16 hour photoperiod at 20°C and an 8 hour dark period at 15°C. Once germinated, all seedlings were moved to a vernalization chamber for 8 weeks with a 10 hour photoperiod and 14 hour dark period at 4°C. Following vernalization, all seedlings were replanted into 1 litre pots in course nutrient-rich compost, transferred to the glasshouse and grown under the same conditions as previously.

Durum Hoh-501 spikes were emasculated between growth stages 55 and 59 on the Zadoks scale^70^ when the spike was between 50% and full emergence from the flag leaf. Less mature florets were removed so that all remaining florets were at a similar developmental stage. Florets were cut to around half their size to allow removal of all immature green anthers. Once anthers had been removed, the emasculated ear was covered with a clear glassine bag to avoid desiccation and cross-pollination. Two or three days post emasculation, *Ae. tauschii* plants with extruding pollen were selected as pollen donors for artificial pollination of receptive Hoh-501 stigmas.

At 3 weeks post pollination, all set seed were removed from the recipient female ear and cleaned in 20% bleach solution for 15 minutes. The embryo of each seed was excised under aseptic conditions and placed on Murashige and Skoog media with 20 g/l sucrose, 2 mg/l zeatin, 2.5 mg/l CuSO4, 6 g/l Sigma type 1 agarose, at pH 5.8. The embryo was placed with the scutellum in contact with the media and the Petri dish sealed with parafilm. Excised embryos were placed on covered trays in a growth room held at 25°C for 1-2 weeks until germination. Embryos that failed to germinate within two weeks were given a cold treatment of 4°C for seven days to encourage germination. Any embryos that failed to germinate after cold treatment were discarded. Germinated plantlets were then vernalized for 8 weeks at 4°C with a 10 hour photoperiod. After vernalization, plantlets were transferred to 1 litre pots containing Levingtons M2 compost and placed in a growth chamber at 20°C with a 16 hour photoperiod and 8 hour dark period at 15°C until they reached growth stage 23 on the Zadoks scale^70^.

At this stage after transplantation the plants were treated with colchicine to double their chromosome complement. A solution of 0.05% colchicine was prepared using 12 ml distilled water, 6 mg colchicine, 0.18 ml dimethyl sulfoxide and a single drop of Tween-20 per plant. Strong tillering (4-6 leaves) haploid seedlings were removed from their pots and excess soil washed from the roots under a running tap. The roots were trimmed to around 2-3 cm and the leaves trimmed to a height of 10-15 cm. The roots of the seedlings were immersed in a beaker containing 13 ml of the pre-prepared 0.05% colchicine solution. A clear plastic bag was placed over the seedlings and two angle-poise lamps were placed either side of the transparent bag to encourage transpiration and therefore the uptake of colchicine through the seedlings. After 5.5 hours of exposure, the seedlings were removed from the colchicine solution and the roots were washed thoroughly under a running tap. The seedlings were potted into moist Levingtons M2 compost and transferred to a growth chamber held at 20°C and a 16 hour photoperiod. The treated seedling will often die back within 1–2 weeks of colchicine treatment and new shoots will emerge shortly afterwards that can be grown to maturity. All emerging ears were bagged and seed of the primary synthetic wheat collected at maturity. The lines are available from the Germplasm Resources Unit www.SeedStor.ac.uk under entry numbers WS0461 to WS0501 (*Ae. tauschii* donor lines), WS0001 to WS0043 (synthetic hexaploid wheat lines) and WS0502 (durum line Hoh-501) (Supplementary Table 11).

Analysis of genetic redundancy based on whole genome shotgun sequencing (see above) subsequently revealed that some of the *Ae. tauschii* donor accessions were genetically redundant with each other. Moreover, we were unable to germinate the seed for harvesting tissue for DNA preps and sequencing for four *Ae. tauschii* donors. Our final set therefore included 32 sequenced and unique non-redundant *Ae. tauschii* donor accessions (Supplementary Table 11).

### Determining interval sizes of designated genes mapped to the D genome using bi-parental genetics

The 2017 Komugi wheat gene index (https://shigen.nig.ac.jp/wheat/komugi/genes/symbolClassList.jsp) was consulted to catalogue designated genes present within the genome of *Ae. tauschii* or the D-subgenome of *T. aestivum*. Candidate genes were filtered based on the existence of the following information within references listed in the Komugi index; (i) the presence of a marker either side of each gene, (ii) the size of the bi-parental population, and (iii) the marker number. The forward and reverse primer sequences for each marker were obtained from GrainGenes (https://wheat.pw.usda.gov/GG3/) or from published literature and BLASTed against the wheat cv. Chinese Spring assembly (IWGSC, INSDC GCA 900519105.1), available at EnsemblPlants. The best candidate location for each marker was used to calculate the mapping interval for each gene (Supplementary Table 15).

## Data availability

The raw PacBio and Illumina sequences used for the assembly of *Ae. tauschii* accession TOWWC0106 have been submitted to the Genome Sequence Archive (GSA) of the National Genomics Data Center hosted by the Beijing Genomics Institute, Beijing, under the accession number CRA002681.

The genome assemblies and annotations of TOWWC0112 and TOWWC0106 are available from the Leibniz Institute of Plant Genetics and Crop Plant Research (IPK) at https://doi.ipk-gatersleben.de/DOI/4bb6f03f-3a15-429a-b542-9962cb676e63/953a2d8a-5ade-479a-9304-6fdd12da7ce4/2/1847940088.

The 150 bp paired-end Illumina sequences for the 306 *Ae. tauschii* accessions, the 250 bp paired-end and mate-pair libraries for accession T0WW0112 and the RNAseq data for eight *Ae. tauschii* accessions is available from NCBI study number PRJNA685125.

The 150 bp paired-end Illumina sequences for the hexaploid wheat accessions and the two additional *Ae. tauschii* accessions used in the *Cmc4* and *Cmc_TAM112_* haplotype analysis (Fig. 4; Supplementary Fig. 16) are available from NCBI study number PRJNA694980.

The *k*-mer matrix for 305 *Ae. tauschii* accessions and the tetraploid donor *T. durum* Hoh-501 used to generate synthetic hexaploids can be obtained from https://doi.ipk-gatersleben.de/DOI/dfc2d351-b5fe-41e6-bd6c-efe96cfcc7aa/0cef0e89-acf2-451c-8efc-a71c0368fec4/2/1847940088.

The variant call (SNP) file for 306 *Ae. tauschii* accessions based on the AL8/78 reference is available from Zenodo under DOI 10.5281/zenodo.4317950.

Counts of lineage-specific *k*-mers in wheat genome assemblies are available from Zenodo under DOI 10.5281/zenodo.4474428.

MEGAHIT assemblies for 303 *Ae. tauschii* accessions (including the 242 non-redundant accessions) are available from Zenodo under DOIs 10.5281/zenodo.4430803, 10.5281/zenodo.4430872 and 10.5281/zenodo.4430891.

A 29,245 bp fragment extracted from contig 00015145 of the *Ae. tauschii* TOWWC0106 assembly was deposited in the NCBI GenBank, along with the coordinates of the *WTK4* transcript SV01, under study number MW295405.

The *SrTA1662* gene and transcript sequence have been deposited in NCBI Genbank under accession number MW526949.

Figures that have associated raw data include Figs. 1-6 and Supplementary Figs. 2-13 and 15-16.

## Code availability

Scripts for SNP calling, *k*-mer matrix generation, redundancy analysis, determination of residual heterogeneity, phylogenetic tree construction including iTOL .nwk files, admixture analysis, *k-*mer GWAS, and SNP GWAS, can be found in the repository https://github.com/wheatgenetics/owwc.

## Acknowledgements

We are grateful to the germplasm banks at Kansas State University Wheat Genetics Resource Center, International Center for Agricultural Research in the Dry Areas, USDA-ARS National Small Grains Collection, Leibniz Institute of Plant Genetics and Crop Plant Research, Ilam University, Tajikistan Academy of Sciences, and the N. I. Vavilov Research Institute of Plant Industry for providing seed and/or collection data of *Ae. tauschii*. We thank our colleagues Y. Yue, P. Crane, S. Burrows and John Innes Centre (JIC) Horticultural Services for plant husbandry, M. Ambrose for help with public distribution of germplasm, M. Craze and S. Bowden for help with creation of synthetic wheats, H. Jones for help with elucidating provenance of *Ae. tauschii* donors used for synthetic wheats, C. Kling for developing and making available the durum wheat line Hoh-501 used for generating synthetic wheats, R. Graf for supplying wheat cv. ‘Radiant’, H. Cherry Guo for managing Illumina sequencing, T. Olsson for Illumina data handling, C. Michael Smith for maintenance of wheat curl mite colonies, H. Ahlers for creating graphics, M. Buttner for helpful discussions, A. Galvin and A. Lawn for OWWC communications, A. Meldrum for drafting OWWC research agreement, the JIC NBI Computing Infrastructure for Science group and the Kansas State University (KSU) BEOCAT for HPC access and maintenance, and S. Krattinger for reviewing the draft manuscript.

This research was financed by the UK Biotechnology and Biological Sciences Research Council (BBSRC) Wheat Improvement Strategic Programme BB/I002561/1 to RH and AB; BBSRC Designing Future Wheat Institute Strategic Programme BB/P016855/1 to RH, AB, PN, SB, XB, RD, CU and BW; BBSSRC Earlham Institute Strategic Programme BBS/E/T/000PR9817 to RD; BBSRC-Embrapa Newton Fund BB/N019113/1 to PN; BBSRC grant BB/PPR1740/1 to WH; BBSRC National Capability award BBS/E/T/000PR9814 to RD; UK Research and Innovation-BBSRC National Capability grant BBS/E/J/000PR8000 to NC; a BBSRC Doctoral Training Partnership scholarship to AH; US National Science Foundation (NSF) Industry-University Cooperative Research Center (IUCRC) Award 1822162 to JP; Phase II IUCRC at KSU Center for Wheat Genetic Resources to JP; US-NSF award Grant/FAIN 1339389 to JP; Kansas Wheat Commission award B65336 to JP; US-NSF award IOS-1238231 to JD and M-CL; United States Department of Agriculture (USDA) to GB-G, SX and JF; the National Institute of Food and Agriculture-USDA to LG; a Fulbright Scholars Program to PS; Swiss National Science Foundation award 310030B_182833 to BK; Newton-Mosharafa Fund award 332408563 to AE and BW; a JIC Institute Development Grant to BW; Agriculture Development Fund of the Saskatchewan Ministry of Agriculture project 20180095 to GB and HRK; Saskatchewan Wheat Development Commission to GB and HRK; Alberta Wheat Development Commission to BG and HRK; Manitoba Crop Alliance to GB and HRK; Government of Saskatchewan Ministry of Agriculture to PH; European Research Council award ERC-2016-STG-716233-MIREDI to KK; a Consejo Nacional de Ciencia y Tecnología scholarship to JQC; JIC International Scholarships to JQC and SG; Monsanto’s (now Bayer) Beachell-Borlaug International Scholars’ Program to SG; 2Blades Foundation to SG and BW; John Innes Foundation to JW; European Union’s Horizon 2020 research and innovation programme Marie Skłodowska-Curie grant agreement 674964 to NK, BW and CU; JIC Science For Africa Initiative to NK; The Royal Society award UF150081 to SB; Australian Research Council award DP210103744 to SB; a Università di Bologna scholarship to AP; Innovation Fund Denmark award 4105-00022B to MP and AJ; Jewish National Fund of Australia to RA and AS; Ministry of Education and Culture of the Republic of Indonesia and the Austrian Agency for International Cooperation in Education and Research (OeAD-GmbH) in cooperation with ASEA-UNINET to RPK; Department of Biotechnology, India award BT/PR30871/BIC/101/1159/2018 to NSa and award BT/IN/Indo-UK/CGAT/14/PC/2014-15 to PC; Science and Technology Development Fund, Egypt-UK Newton-Mosharafa Institutional Links award 30718 to A.E. and B.W. National Science Foundation of China grants 91731305 and 31661143007 to LM; Knowledge Innovation Program of Chinese Academy of Agricultural Sciences award CAAS- DRW202002 to LM; the breeding companies KWS, Limagrain, Syngenta and Bayer to the Open Wild Wheat Consortium.

## Author information

### Contributions

Configured, bulked, and/or distributed *Ae. tauschii* germplasm (S.Aro., M.F., C.G., N.Si., J.Ra., N.C., S.G., A.H., T.O., J.L.). Collected and curated new *Ae. tauschii* accessions from Iran (A.M.) and Tajikistan (F.N.). Extracted plant DNA (M.F., S.W., J.L., A.P., S.Aro., G.Y.) and RNA (A.E., S.Aro.). Prepared DNA libraries (S.W.). Acquired DNA sequences (B.W., J.P., J.F., G.B.-G., S.X., P.C., K.K., A.S., E.L., J.D., M.-C.L., K.M., A.B., B.J.S., V.T.). Undertook sequence data curation, back-up and/or distribution (K.G., J.Ch., M.M., S.Aro., B.Steu., L.G., ST., X.B., R.D., M.Si., L.S.). Performed variant calling and filtering (L.G., K.G.), *Ae. tauschii* redundancy analysis (L.G., S.Aro., K.G.) and heterogeneity analysis (K.G.). Assembled genomes of TOWW0112 (K.G., M.M.), TOWWC0106 (A.L., L.M., D.L.), and diversity panel (K.G.). Performed genome annotations (T.L., S.Art., K.M.). Performed genome-wide phylogenetic analysis (K.G., S.Aro., J.Ch.). Characterized L3 and discovered its contribution to wheat (K.G.). Performed F_ST_ analysis (S.Aro.). Phenotyped *Ae. tauschii* accessions for stem rust (N.K., S.Aro., O.M., B.J.S.), flowering time (J.Q.C., J.S., C.U., B.Stei., R.P.K., H.B.), trichomes (C.C., S.P., P.N.), spikelets (G.B., H.R.K., P.H.), powdery mildew (J.S.-M.) and wheat curl mite (P.S.). Established *k*-mer GWAS methodology and discovered candidate genes (KG). Performed genome-wide LD analysis (K.G., S.Aro.). Performed GWAS control experiments (S.K., K.G., L.G.). Interpreted *Ae. tauschii* trait-genotype relationships for flowering time (J.Q.C., C.U.), trichomes (S.P., S.Aro.), spikelets (S.B., J.W., K.G., J.L.), powdery mildew (J.S.-M., S.Aro., K.G.) and wheat curl mite (P.S., L.G., S.Aro.). Determined gene level (K.G.) and haplotype (K.G., P.S., S.Aro., L.G.) distribution in *Ae. tauschii* and wheat. Estimated genetic diversity captured by wheat landraces and synthetic wheats (K.G.). Annotated *WTK4* (S.Aro., J.S.-M.). Determined *WTK4* gene structure and/or performed functional analysis (J.S.-M., V.W.). Generated synthetic wheats (R.H., A.B.). Developed wheat germplasm for curl mite resistance (S.L., J.Ru.). Annotated *SrTA1662* (S.Aro.), designed and engineered binary construct (M.A.S., S.Aro.), transformed wheat (S.H., W.H.), and phenotyped transgenics (N.K., M.P., A.J., S.Aro.). Designed figures (S.Aro., K.G., J.S.-M., P.S., C.G., T.L., B.W., J.P.). Conceived and designed experiments (K.G., S.Aro., P.S., J.S.-M., L.G., G.B., C.C., C.U., M.M., A.B., B.K., J.P., B.W.). Drafted manuscript (B.W., K.G., P.S., J.S.-M., R.H., S.Aro., J.P., D.G., R.A., L.G., C.G., N.Sa., A.P., S.H., M.S., M.P., C.U., M.M., B.K., K.M., A.S.). Conceived, founded and/or managed OWWC (B.W., J.P., B.Steu.). All authors read and approved the manuscript.

### Ethics declaration

K.G. and B.W. are inventors on UK patent application PC931335GB, which is based on part of the work presented here.

## Supplementary Figures

**Supplementary Fig. 1.**
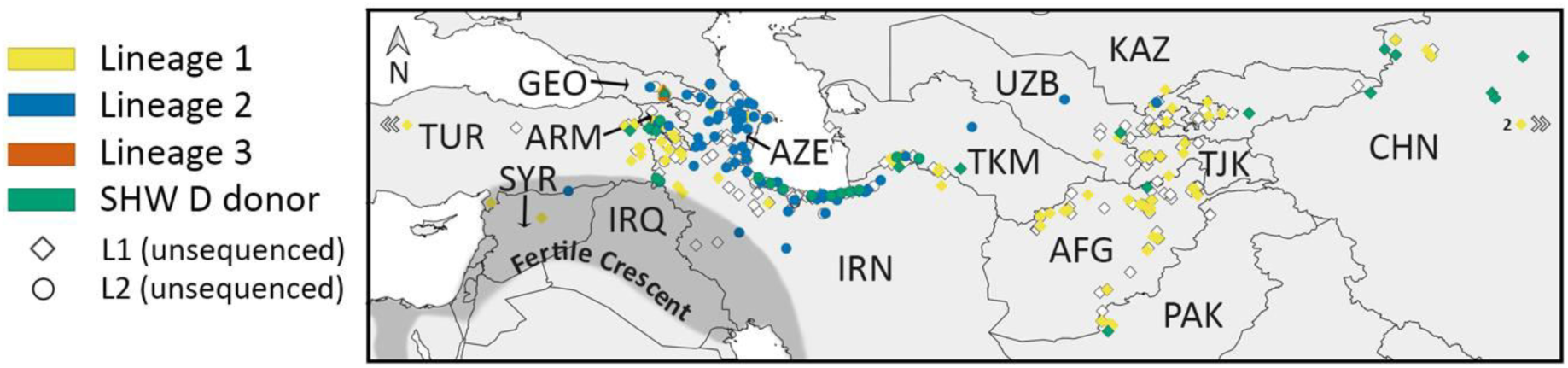
Geographical distribution of *Ae. tauschii* accessions. Filled squares and circles represent accessions sequenced as part of this study, while accessions represented by unfilled squares and circles were not sequenced. Accessions highlighted in yellow were used as D genome donors to generate synthetic hexaploid wheat (SHW) lines. Three accessions outside of the map, one from Turkey and two from China, are indicated by white arrow heads.

**Supplementary Fig. 2.**
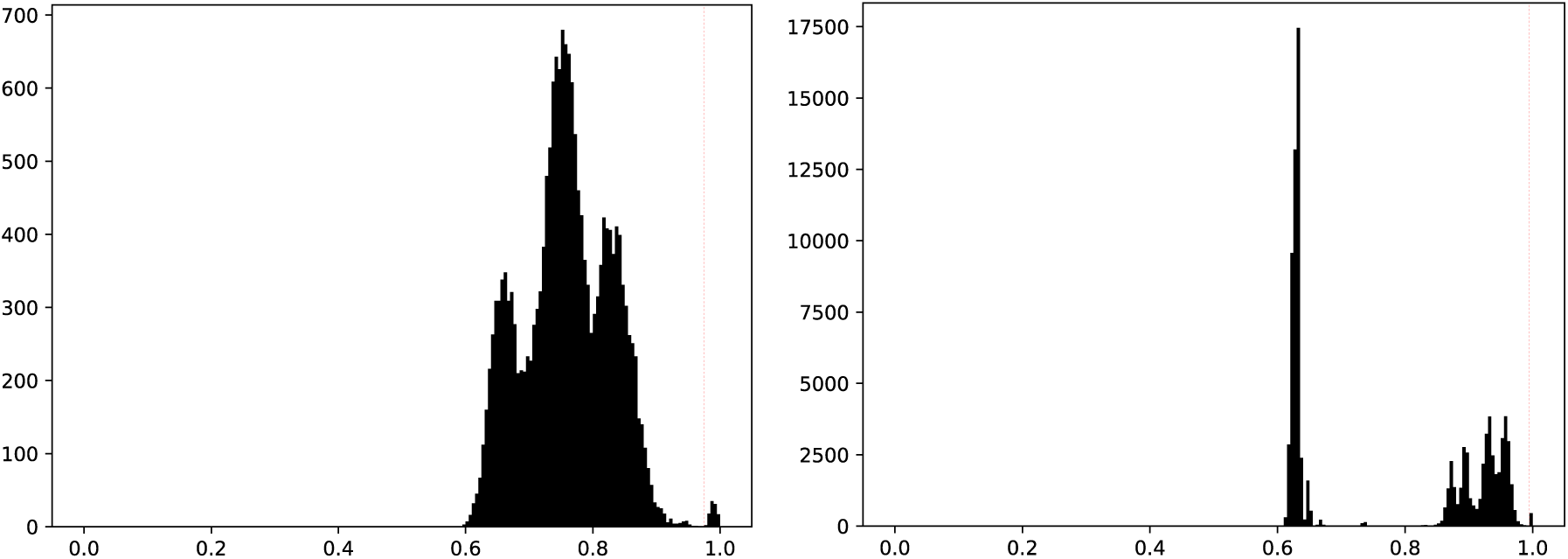
Identification of non-redundant *Ae. tauschii* accessions. Histograms of similarity scores obtained using KASP markers on 195 accessions (left) and 100,000 random SNPs obtained from whole genome shotgun sequencing of 306 accessions (right). The vertical red line in both plots indicates the redundancy cut-off at which the peak of the high similarity values is clearly separated from the rest.

**Supplementary Fig. 3.**
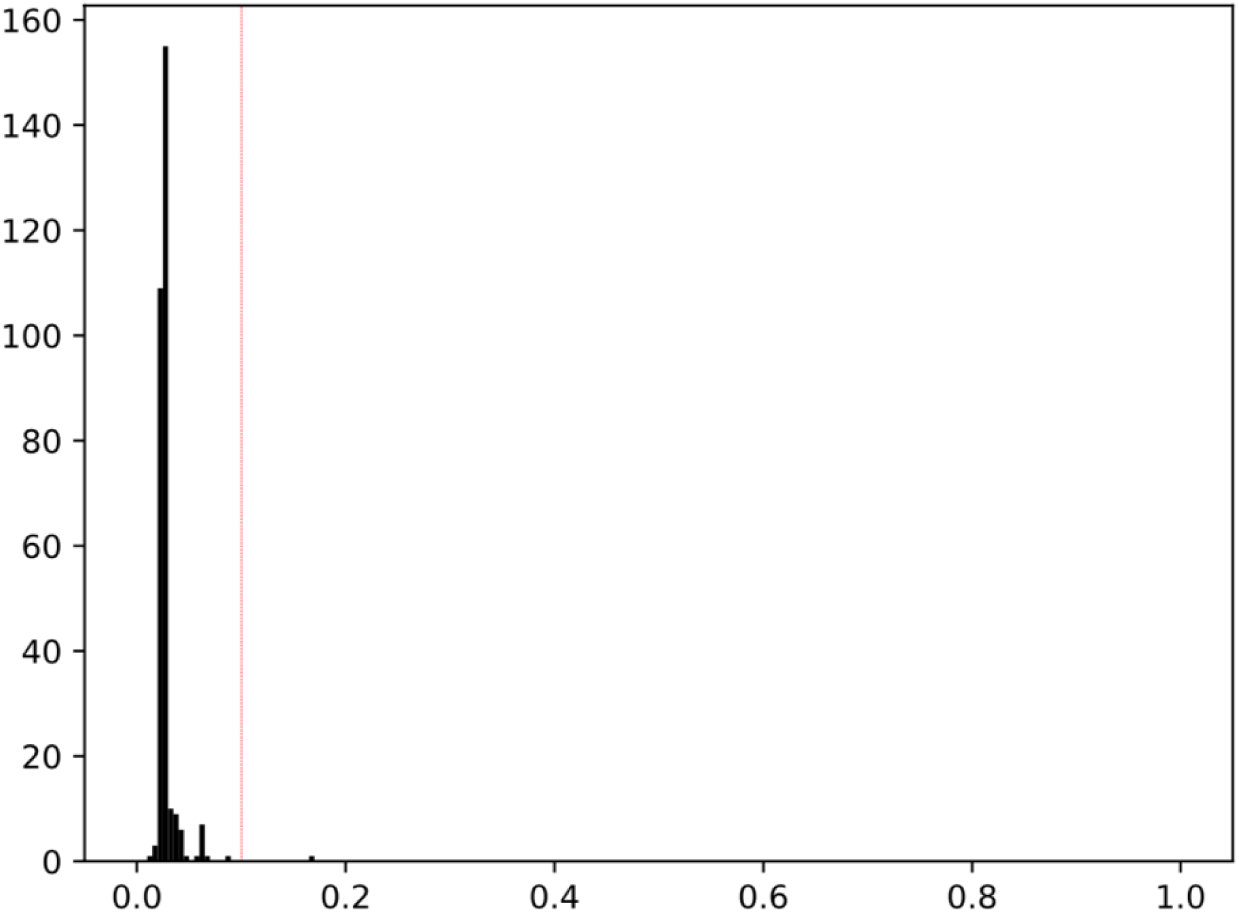
Identification of *Ae. tauschii* accessions with minimal residual heterogeneity. The histogram of heterozygosity scores was generated using all the bi-allelic SNPs obtained from whole genome shotgun sequencing of 305 accessions (excluding TOWWC0193). The vertical red line indicates the cut-off at which the cluster of the low heterozygosity values is clearly separated.

**Supplementary Fig. 4.**
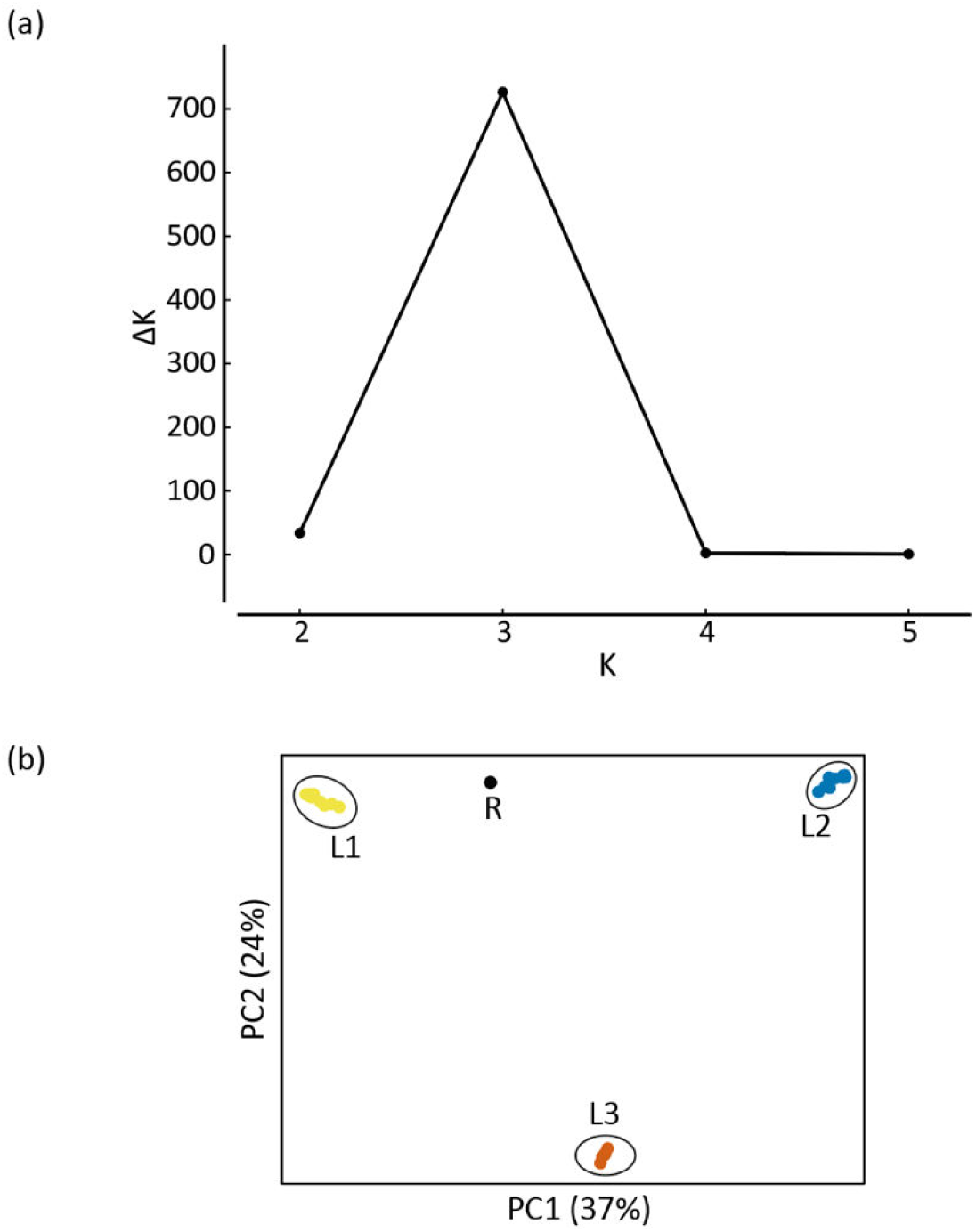
**a**, ΔK plot for a STRUCTURE run with 10 randomly selected accessions each of L1 and L2 along with the five accessions of the putative L3 and the control L1-L2 RIL. **b**, Principal Component Analysis with the same set of accessions as used in panel a. The recombinant inbred control line is indicated by R.

**Supplementary Fig. 5.**
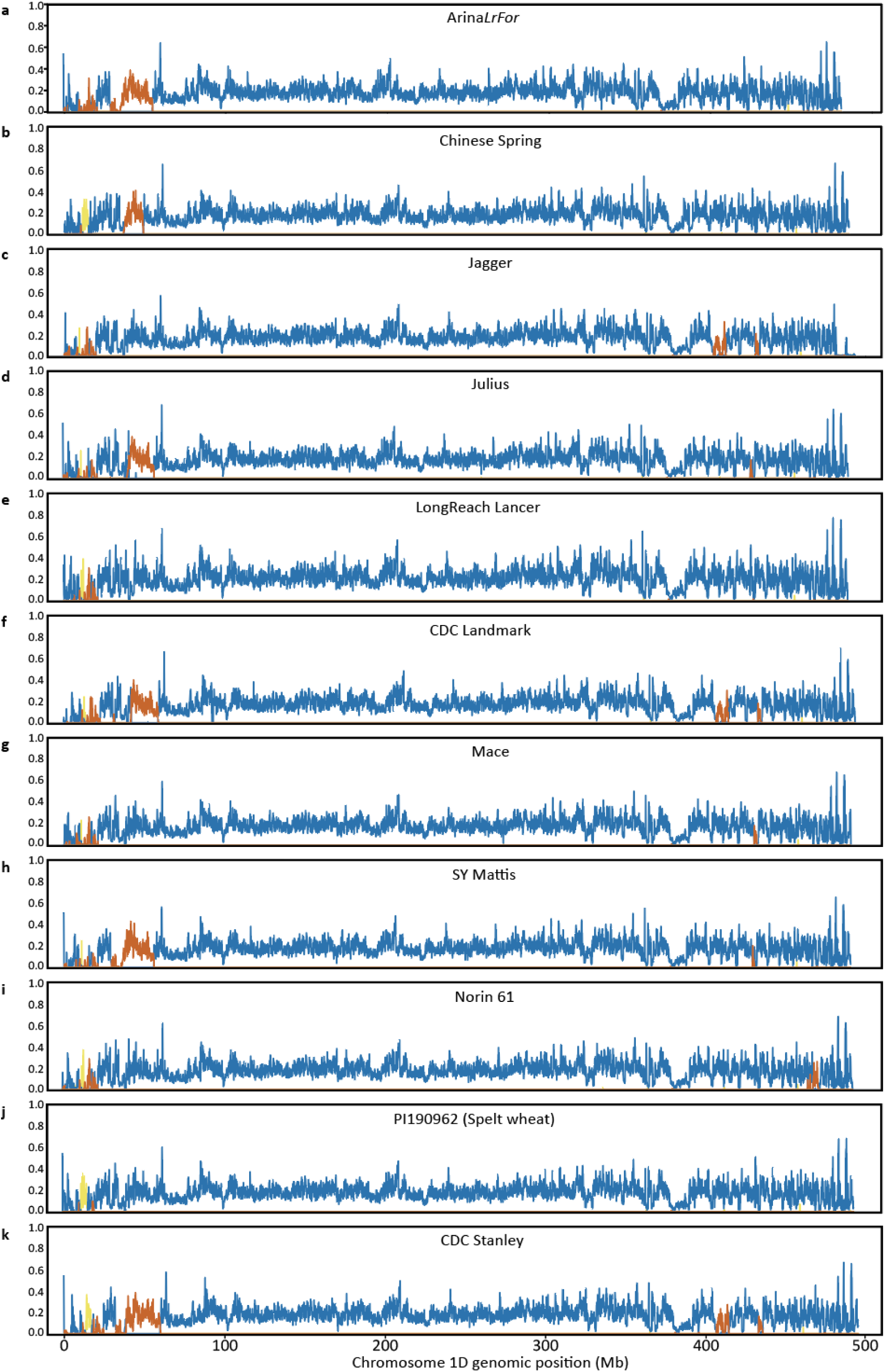
Fraction of lineage-specific *k*-mers in non-overlapping 100 kb windows of Chromosome 1D for the 11 wheat genome assemblies^21^. For the nine modern cultivars, only those *k*-mers were considered which were also present in the short-read sequences of 28 hexaploid wheat landraces^17^.

**Supplementary Fig. 6.**
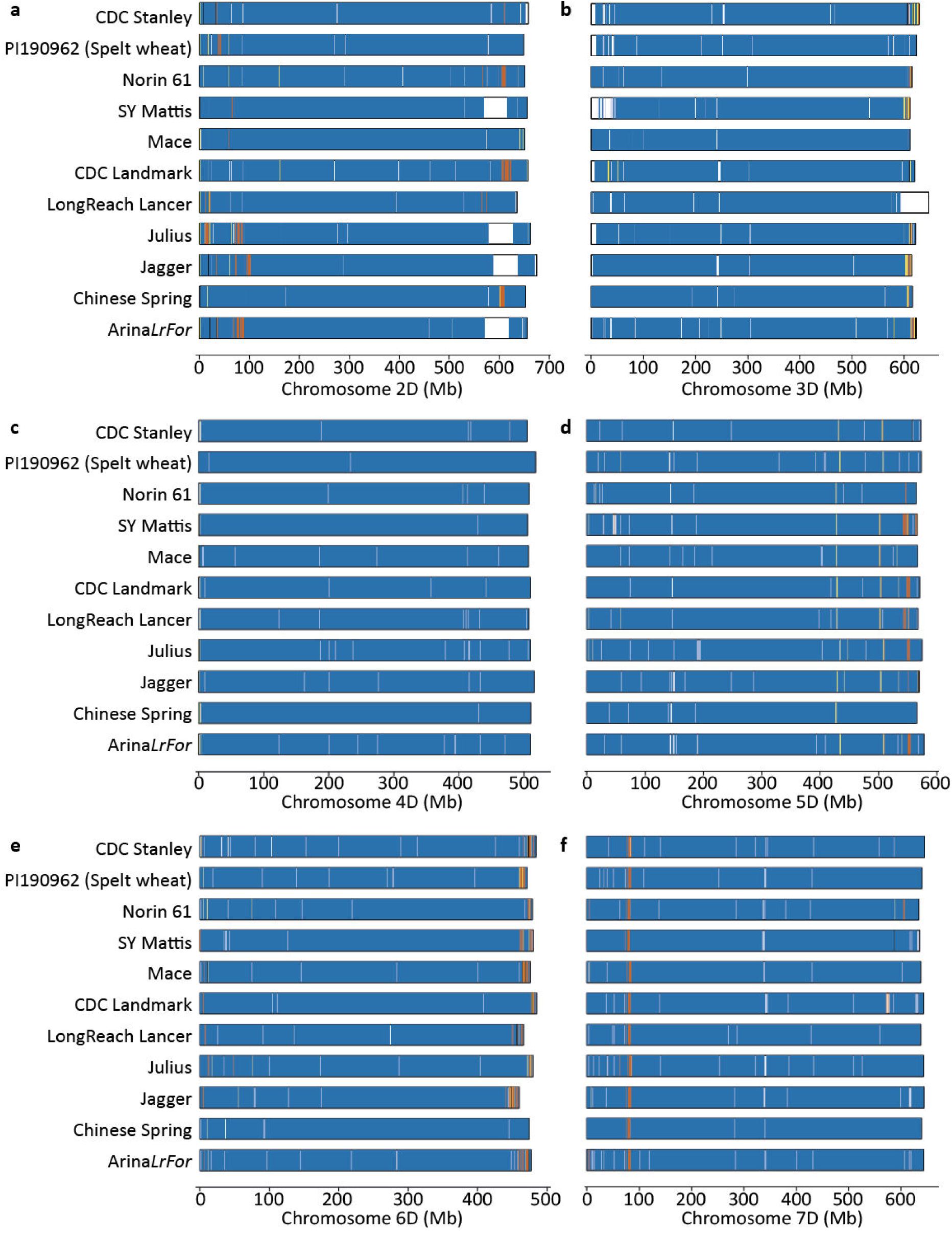
Chromosomes 2D-7D of 11 wheat cultivars colored according to their *Ae. tauschii* lineage-specific origin as in Fig. 1f.

**Supplementary Fig. 7.**
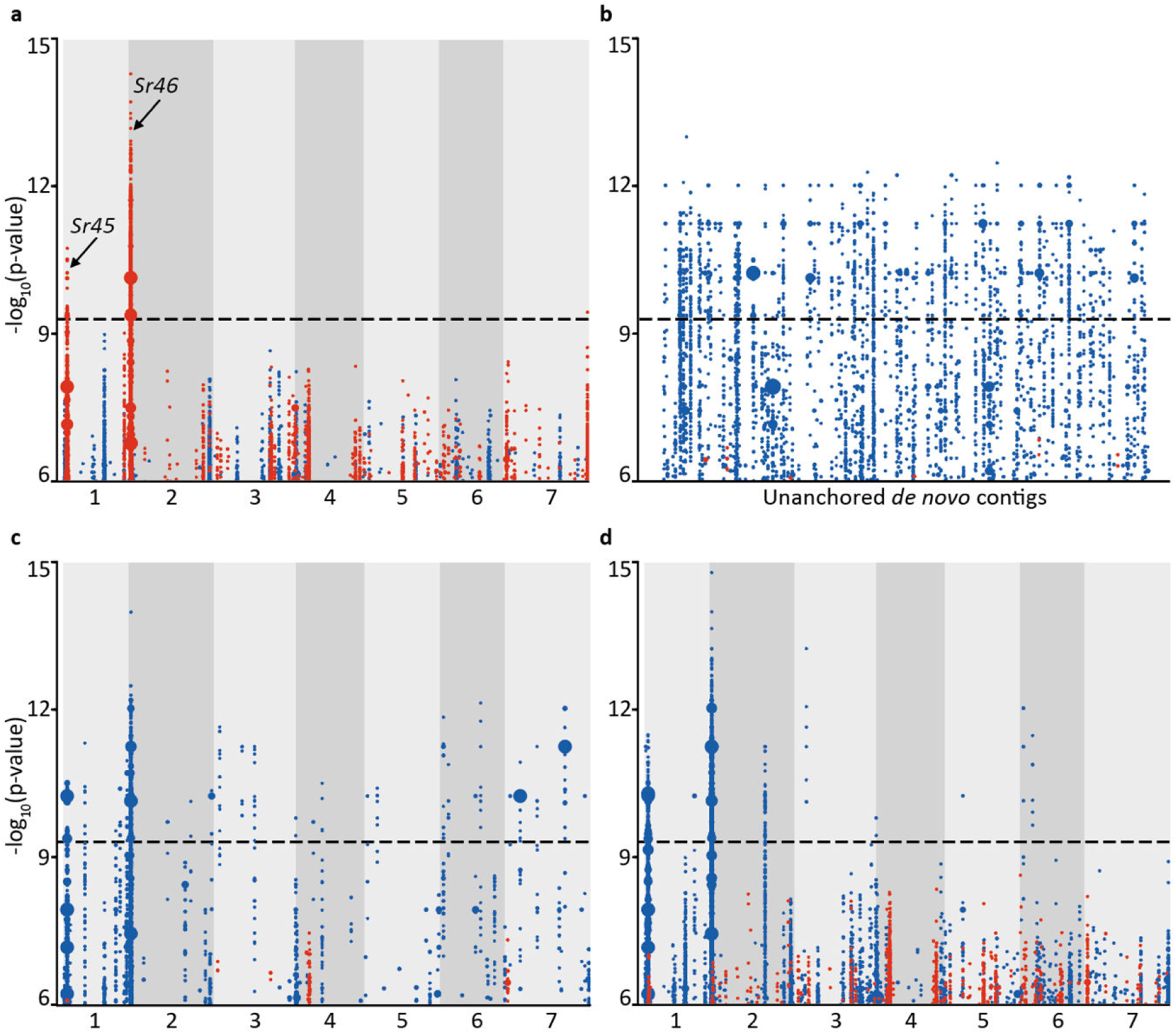
Optimization of *k*-mer GWAS with the positive controls *Sr45* and *Sr46*. Blue/red dots on the y-axis represent one or more *k*-mers significantly associated with resistance/susceptibility, respectively, to *Puccinia graminis* f. sp. *tritici* isolate 04KEN156/04 (race TTKSK) across the diversity panel. Dot size is proportional to the number of *k*-mers having the specific value on the y-axis. **a**, Significantly associated *k*-mers mapped to AL8/78 which is susceptible to TTKSK. The peaks marked *Sr45* and *Sr46* contain the non-functional (not providing resistance to TTKSK) alleles of *Sr45* and *Sr46*. The x-axis represents the seven chromosomes of *Ae. tauschii* reference accession, AL8/78. Each dot column represents a 10 kb interval. **b**, Significantly associated *k*-mers mapped to the unordered *de novo* assembly of TOWWC0112 (N50 1.1 kb), an *Ae. tauschii* accession resistant to TTKSK. Each dot-column on the x-axis represents an unordered contig from the de novo assembly. **c**, Significantly associated *k*-mers mapped to the same assembly of TOWWC0112 as in (b), but now each contig has been ordered by anchoring to the reference genome of AL8/78 (x-axis). **d**, Association mapping with an improved TOWWC0112 assembly (N50 196 kb) anchored to the AL8/78 reference genome (x-axis).

**Supplementary Fig. 8.**
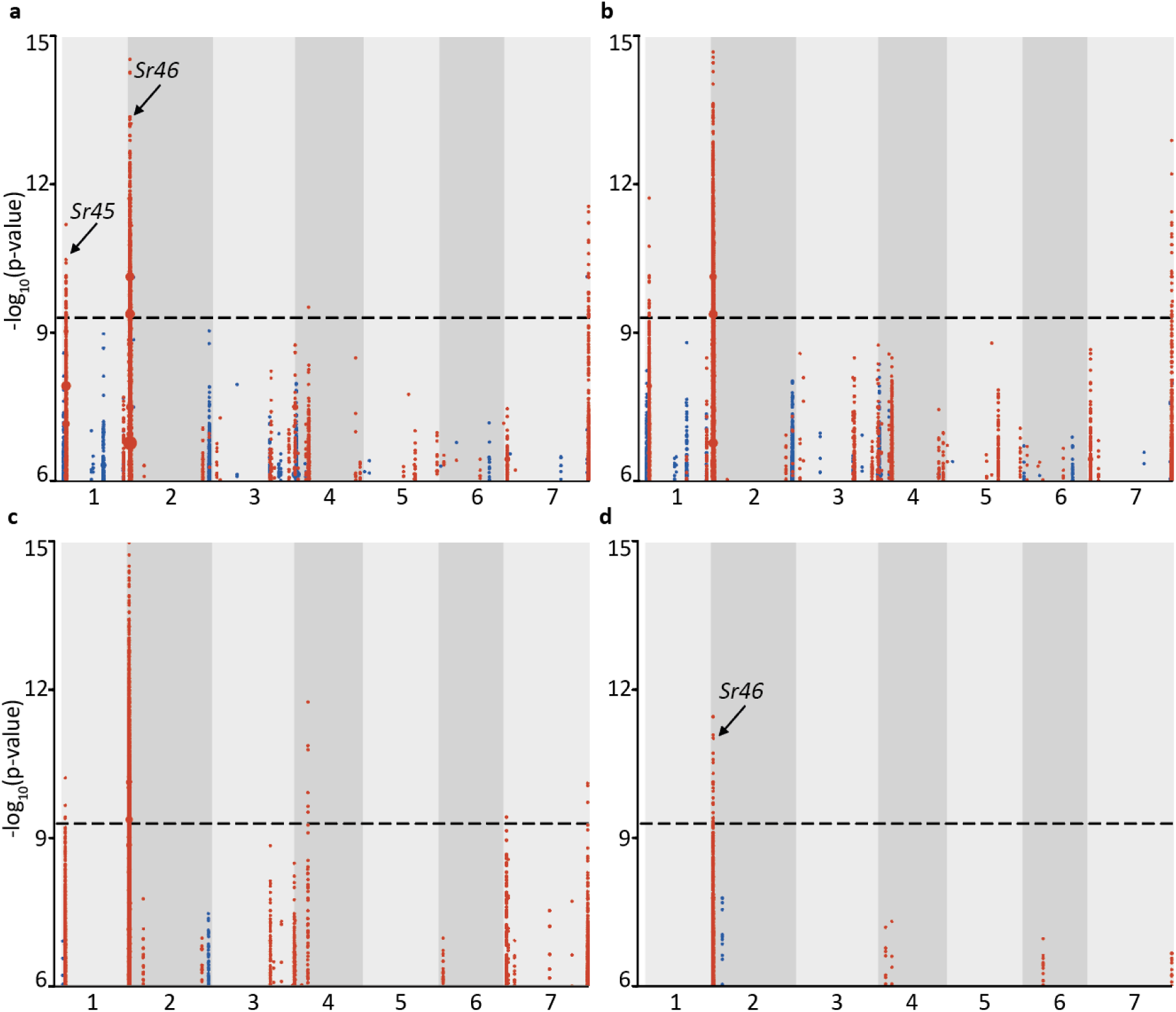
Impact of sequencing coverage on the power to detect the positive controls, *Sr45* and *Sr46.* Sequencing coverage was artificially reduced by sub- sampling the original 10-fold coverage sequencing reads and mapping associated *k*-mers to AL8/78. **a**, Plot obtained with 7.5-fold coverage (compare with 10-fold coverage in Supplementary Fig. 7a). **b**, Plot obtained with 5-fold coverage. **c**, Plot obtained with 3-fold coverage. **d**, Plot obtained with 1-fold coverage.

**Supplementary Figure 9.**
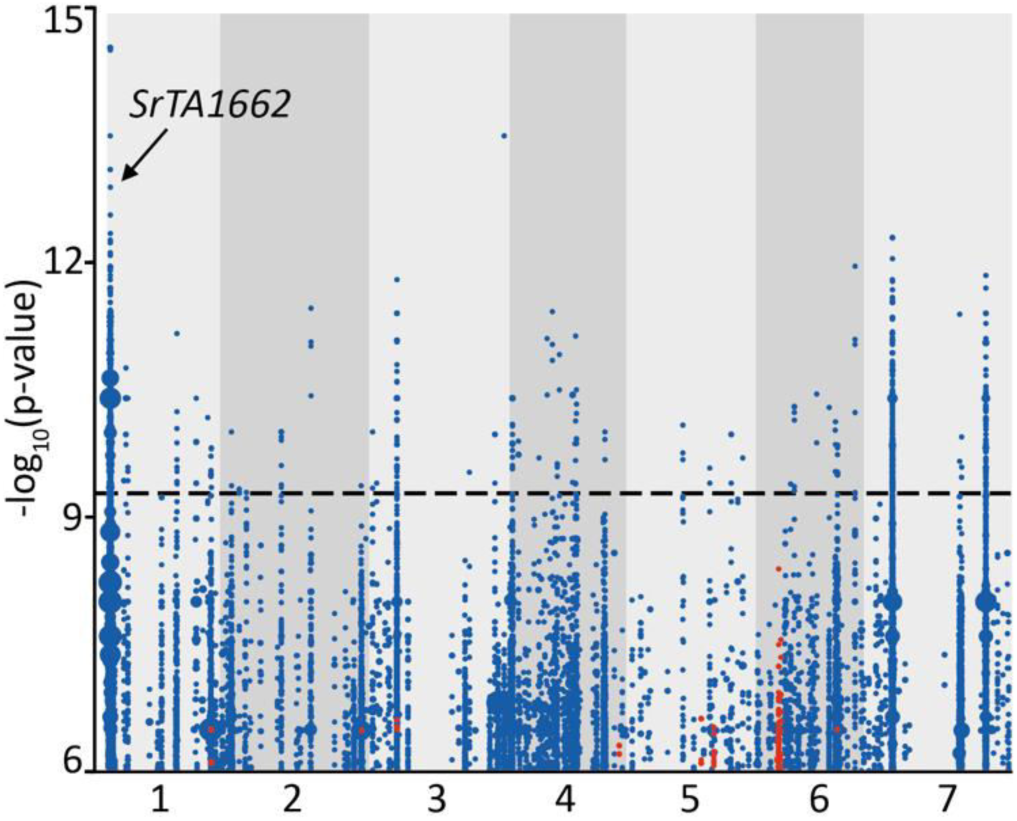
Resistance to *Puccinia graminis* f. sp. *tritici* isolate UK-01 maps to the *SrTA1662* locus. The peak indicated by the arrow contains the region delimited by the *SrTA1662* LD block obtained with *P. graminis* f. sp. *tritici* race QTHJC.

**Supplementary Fig. 10.**
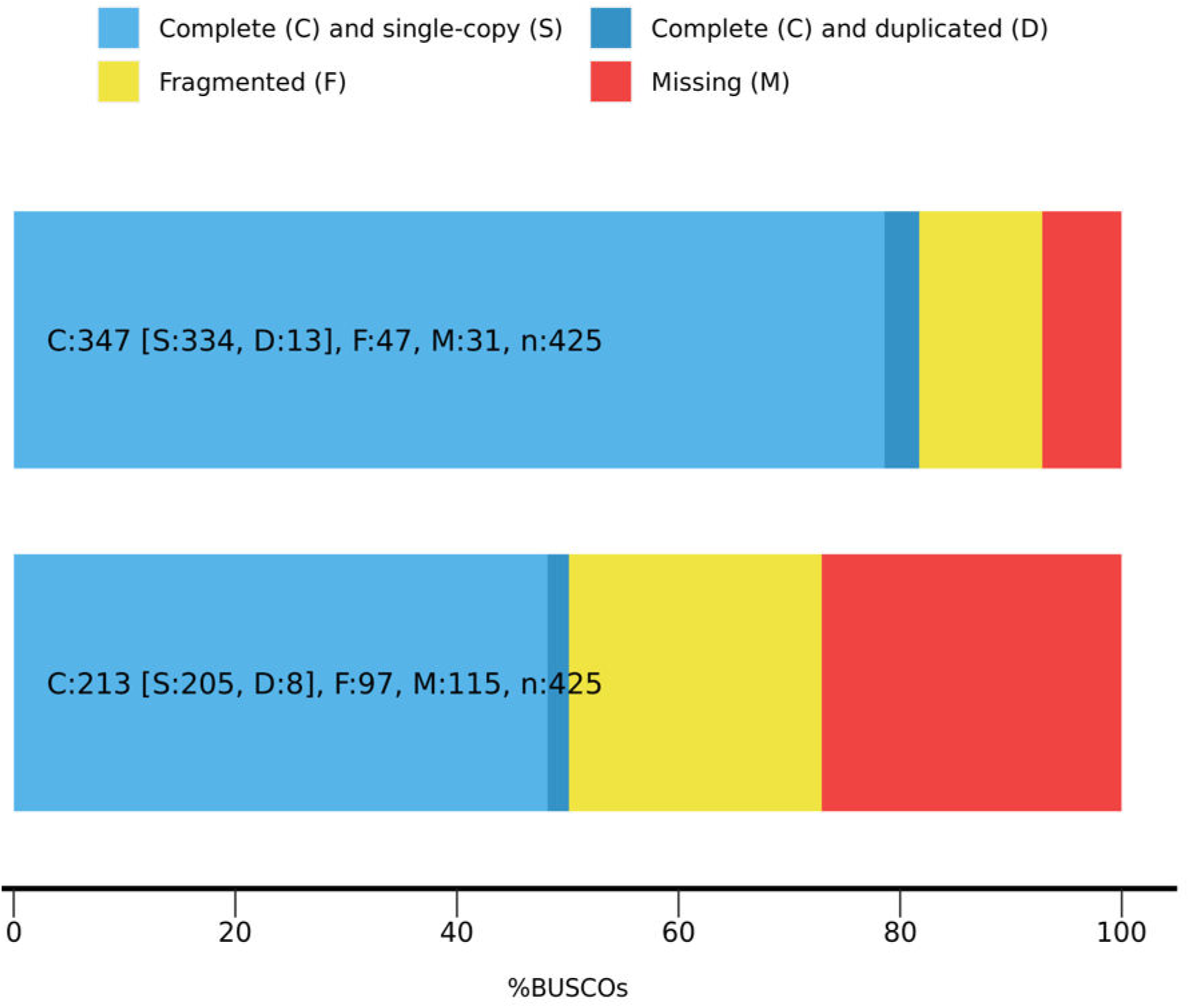
BUSCO assessment results of the annotations of the genome assemblies of the *Aegilops tauschii* accessions TOWWC0106 (top) and TOWWC0112 (bottom).

**Supplementary Fig. 11.**
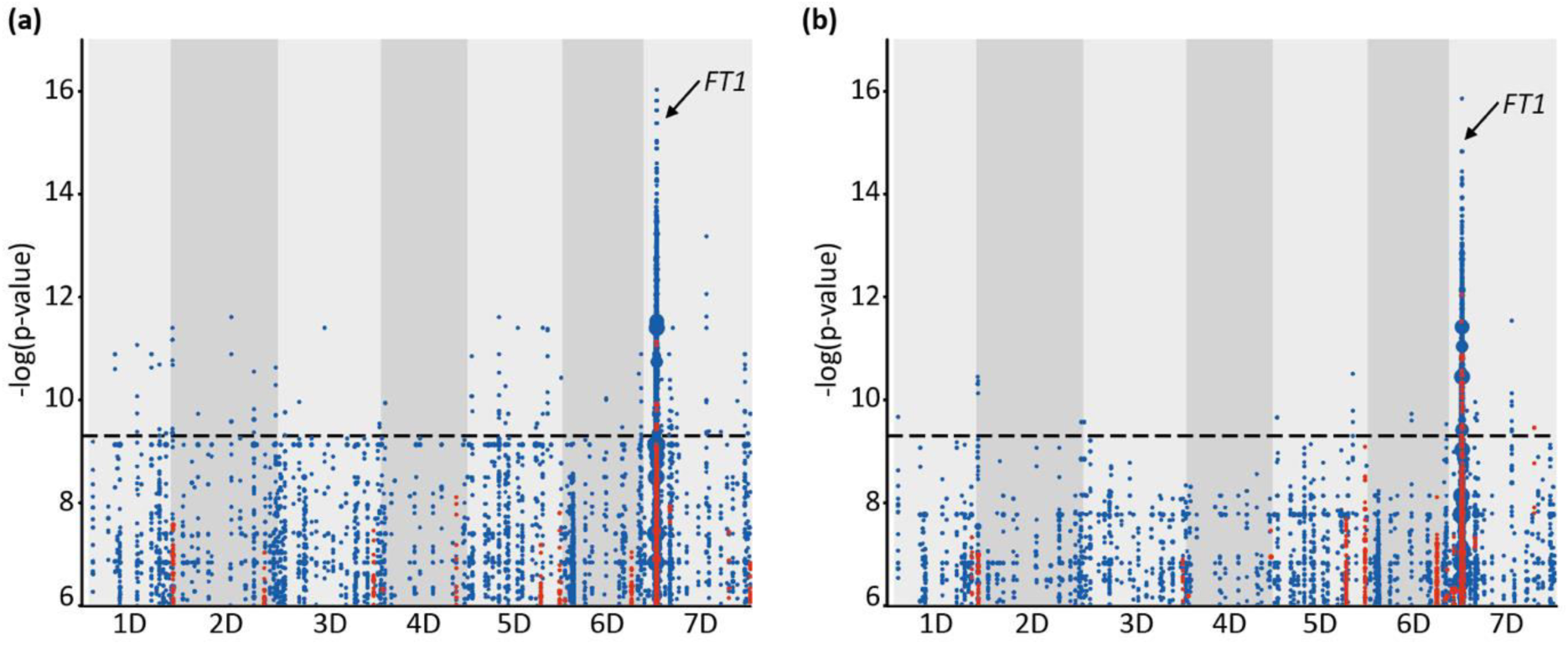
Two independent biological replicates for flowering time identify *k*-mers significantly with *FLOWERING LOCUS T*. The associated *k*-mers were mapped to the *Aegilops tauschii* AL8/78 reference genome where they define a peak similar to that in Figure 2b. **a**, Biological replicate 2. **b**, Biological replicate 3.

**Supplementary Fig. 12.**
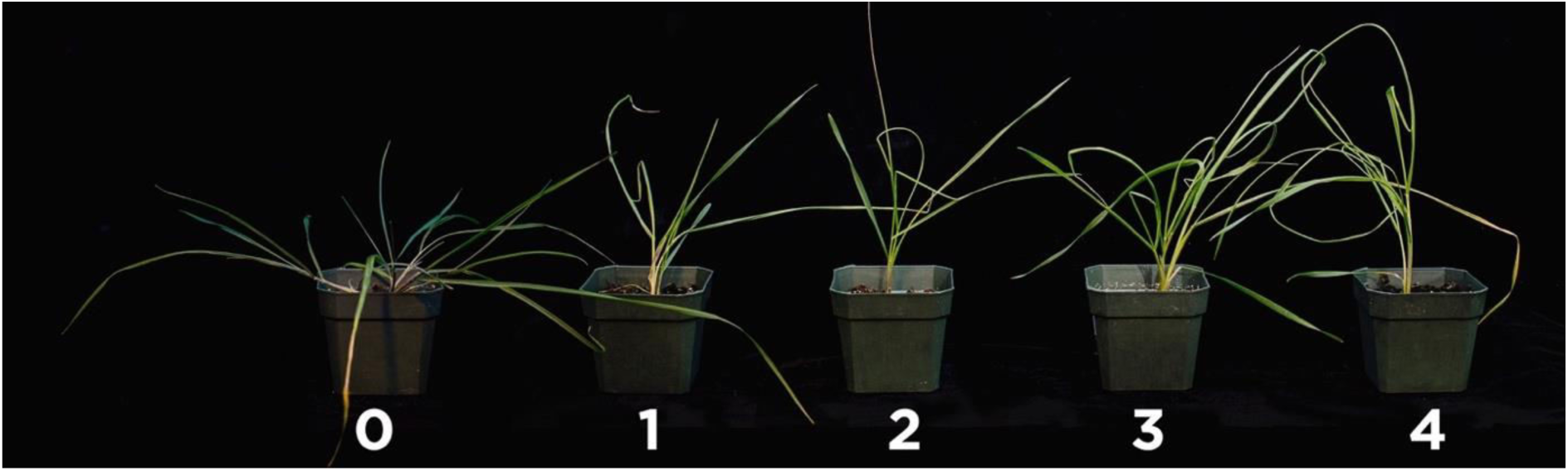
Wheat curl mite (WCM) symptoms and phenotyping scale. Scale used to characterize *Aegilops tauschii* response to WCM infestation. Symptoms used were leaf trapping and leaf curliness. The visual scale ranged from 0 to 4, with 0 equivalent to no symptoms and 1 to 4 denoting increasing levels of curliness or trapped leaves indicative of susceptibility.

**Supplementary Fig. 13.**
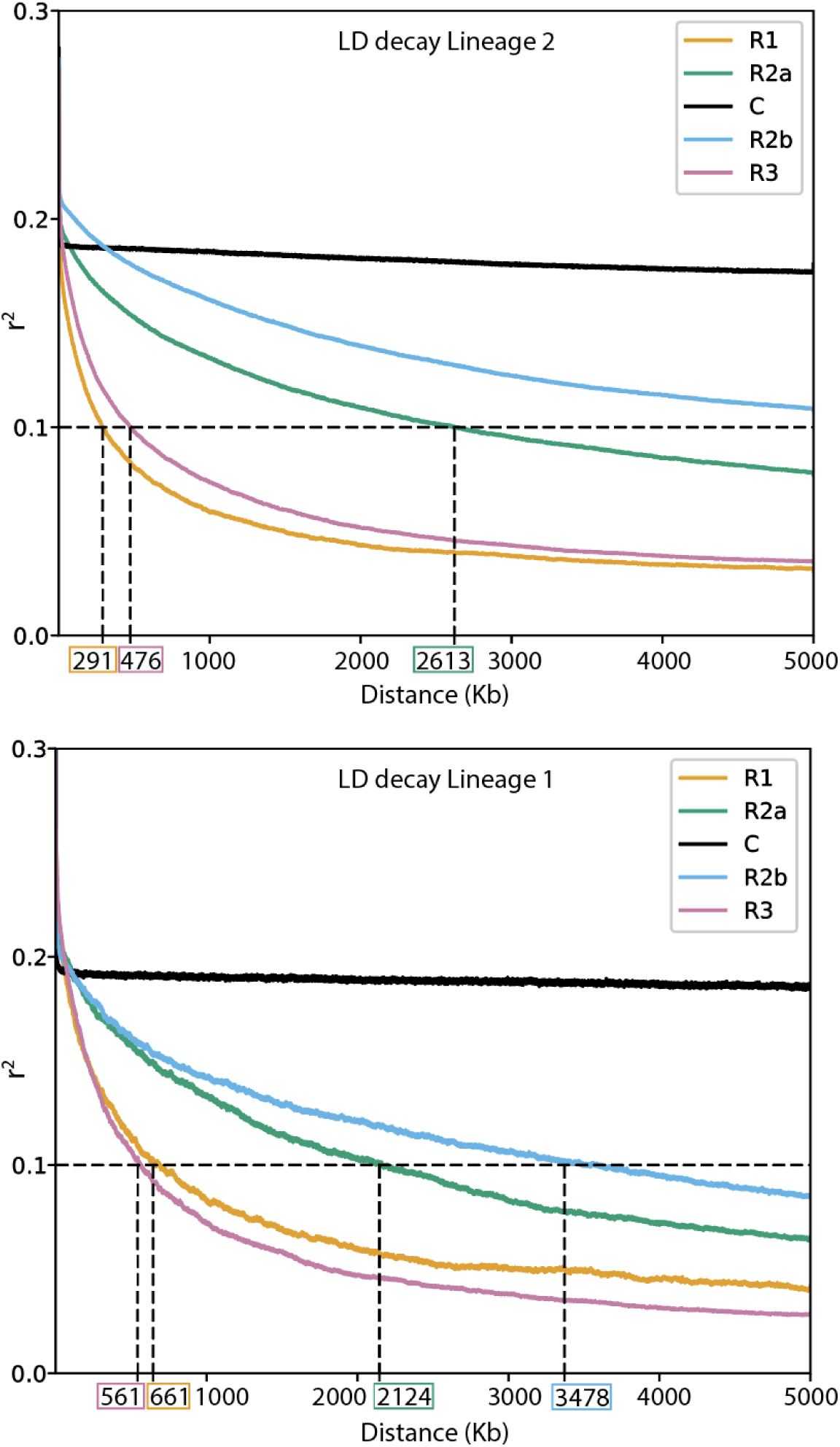
Genome-wide decay of linkage disequilibrium (LD) in *Ae. tauschii*. Genomic regions (R1, R2a, C, R2b, R3) in L2 (top) and L1 (bottom) were determined based on the distribution of the recombination rate in *T. aestivum* cv. Chinese Spring. The distance at which r^2^ for a region drops below 0.1 is highlighted.

**Supplementary Figure 14.**
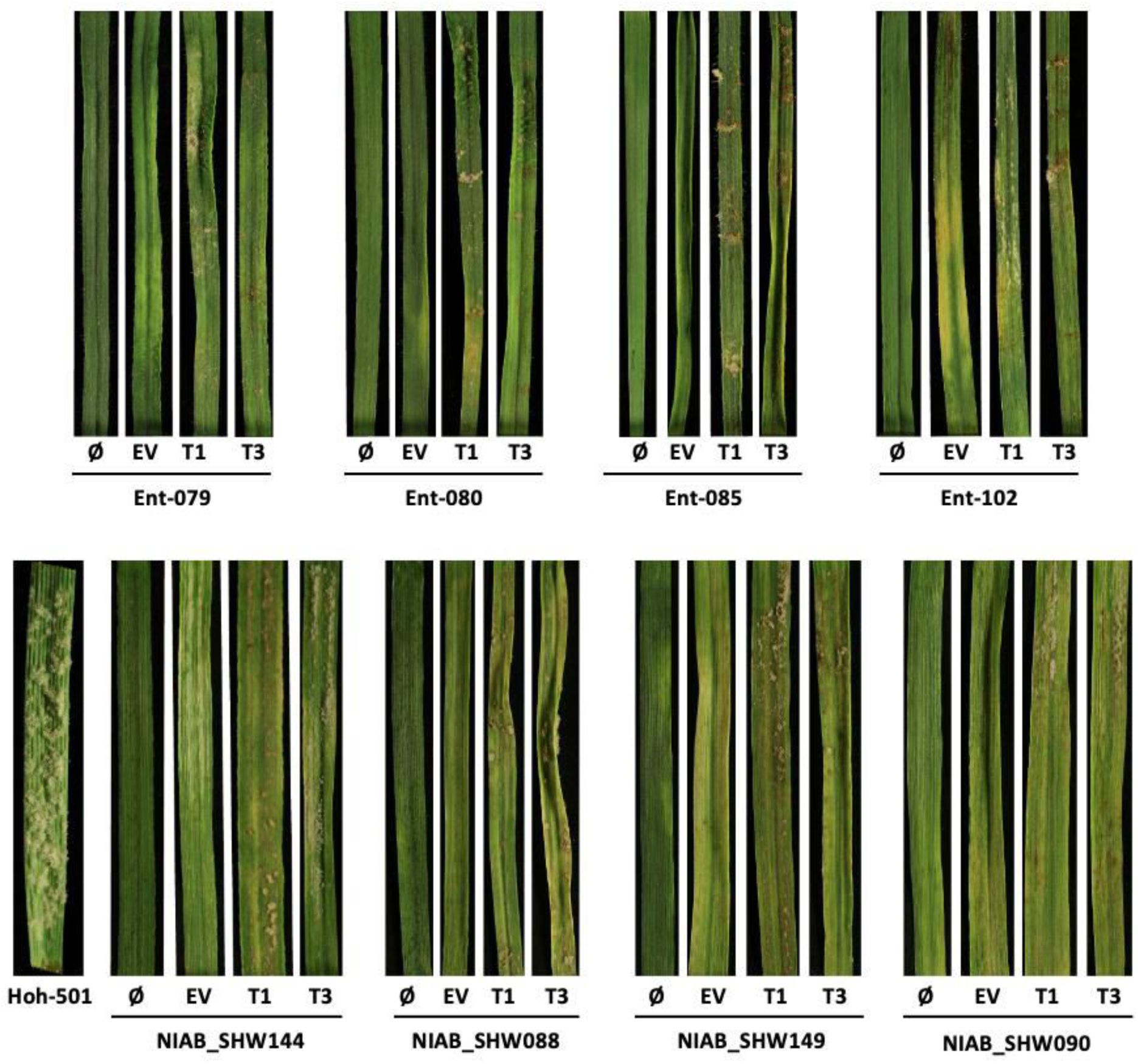
Analysis of powdery mildew resistance in *Aegilops tauschii* and durum donors and their derived synthetic hexaploid wheat lines. . Top, disease reactions to *Blumeria graminis* f. sp. *tritici Bgt96224* are displayed for the *Ae. tauchii* accessions Ent-079, Ent-080, Ent-085 and Ent-102. Bottom, disease reactions to *Bgt96224* are displayed for the corresponding synthetic hexaploid lines (NIAB_144, derived from Ent- 079; NIAB_088 derived from Ent-080; NIAB_149 derived from Ent-085; and NIAB_090 derived from Ent-102) using the tetraploid durum wheat donor line Hoh-501, which is highly susceptible to *Bgt96224.* Each *Ae. tauschii* and its corresponding synthetic hexaploid line was not inoculated with BSMV (Ø) or with a BSMV construct as empty vector (EV) or targeting for silencing the *WTK4* exon 8 (target 1, T1) or exon 10 (target 2, T2), respectively, and then super-infected with *Bgt96224*.

**Supplementary Figure 15.**
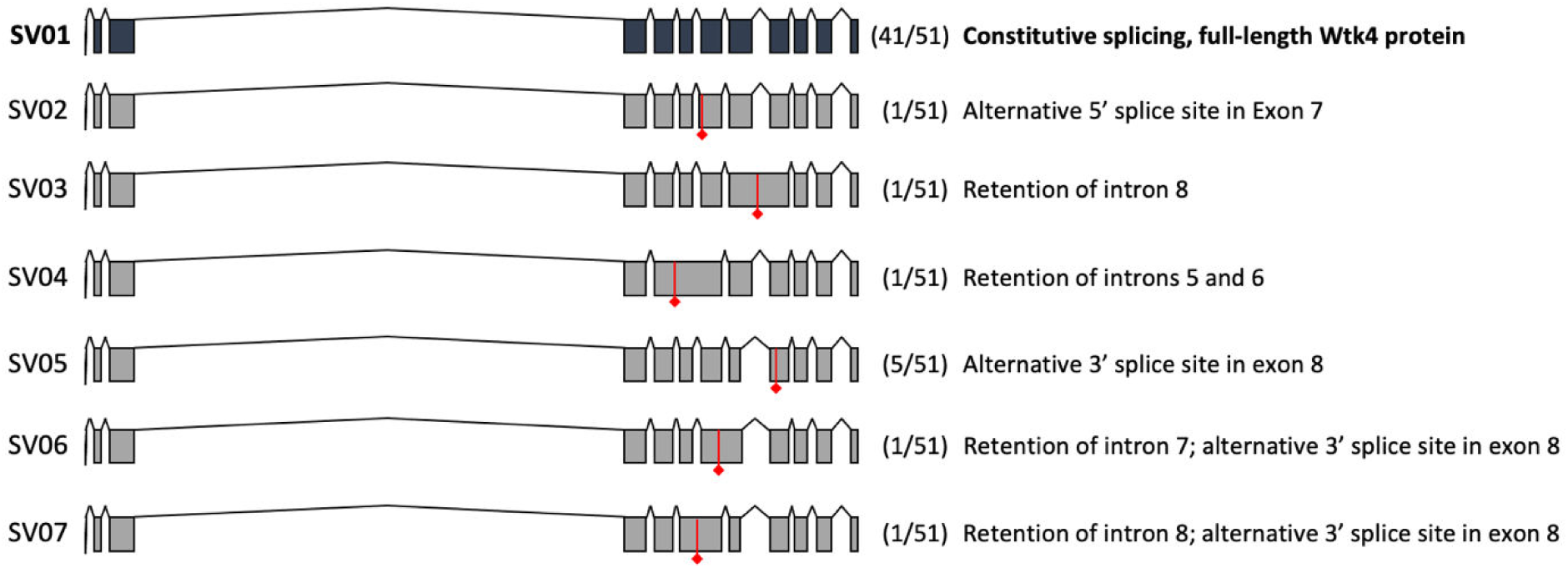
Alternative splicing of *WTK4*. Alternative splicing variants (SV1-7) revealed by sequencing 51 *WTK4* cDNAs. At the top, in black, is shown the splicing variant SV01, which encodes the complete WTK4 protein. Below SV01, six aberrant alternative splicing variants (SV02 to SV07) are shown in in grey. The number of clones identified for each SV is identified in parenthesis. Diamond arrowed red lines point to the first stops codons at the protein level.

**Supplementary Fig. 16.**
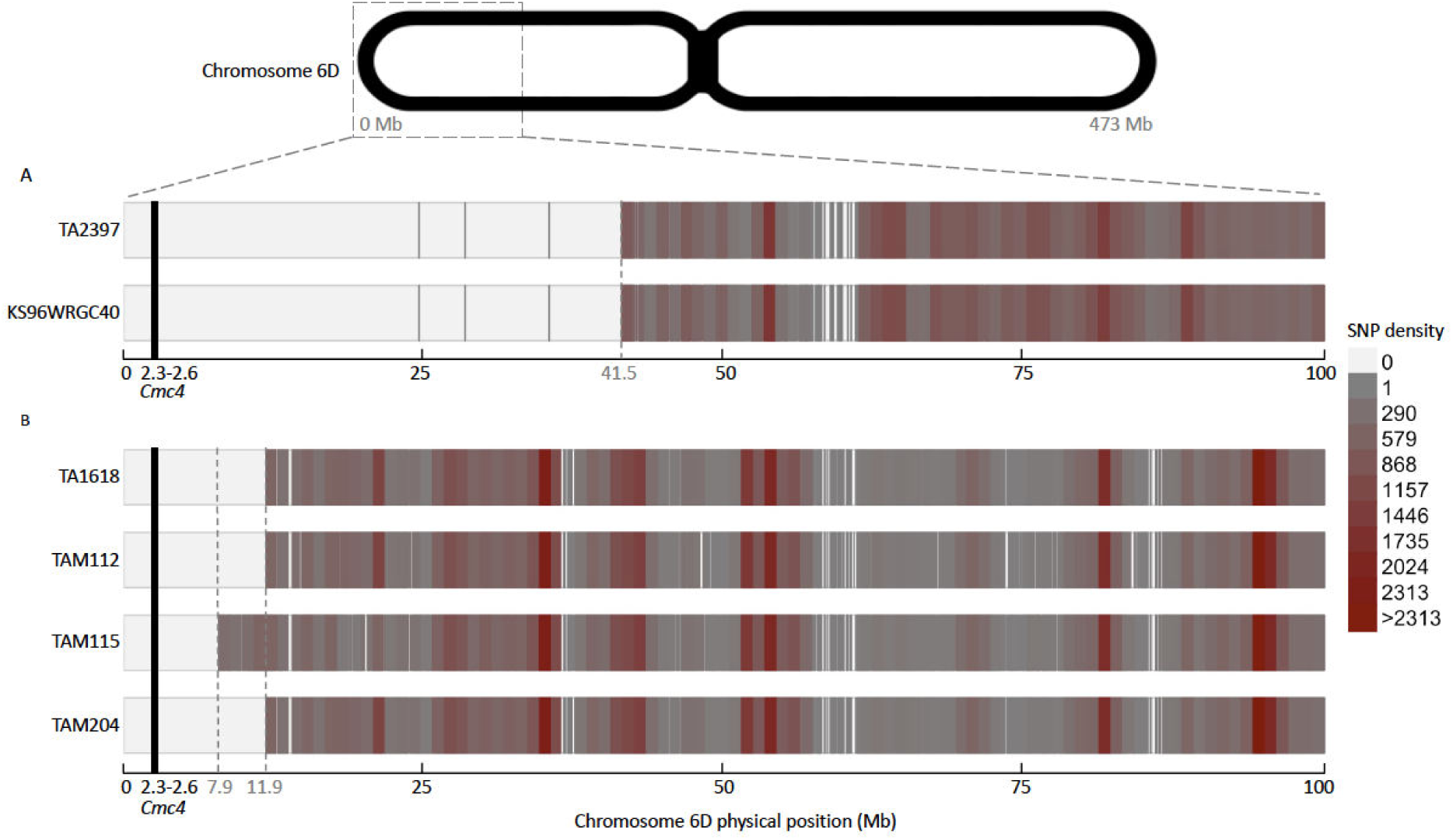
Delineation of *Aegilops tauschii* introgressions carrying wheat curl mite resistance in four wheat lines. The retained polymorphic markers were obtained by pairwise comparisons of the *Ae. tauschii* donor with the corresponding wheat line. **a**, The donor of wheat curl mite resistance in wheat line ‘KS96WGRC40’ is Lineage 1 accession TA2397. This is also the original line where *Cmc4* was mapped. **b**, The donor of resistance in wheat line ‘TAM 112’ is the Lineage 2 accession TA1618. Wheat lines ‘TAM 115’ and ‘TAM 204’ are both resistant through TAM 112. The black vertical line indicates the *Cmc4* position. The three grey dashed vertical lines denote the size of the introgressed fragments, 7.9 Mb, 11.9 Mb, and 41.5 Mb, in the wheat lines ‘TAM 115’, ‘TAM 112’ and ‘TAM 204’, and ‘KS96WGRC40’, respectively. SNP density is based on number of SNPs within 1 Mb bins.

**Supplementary Fig. 17.**
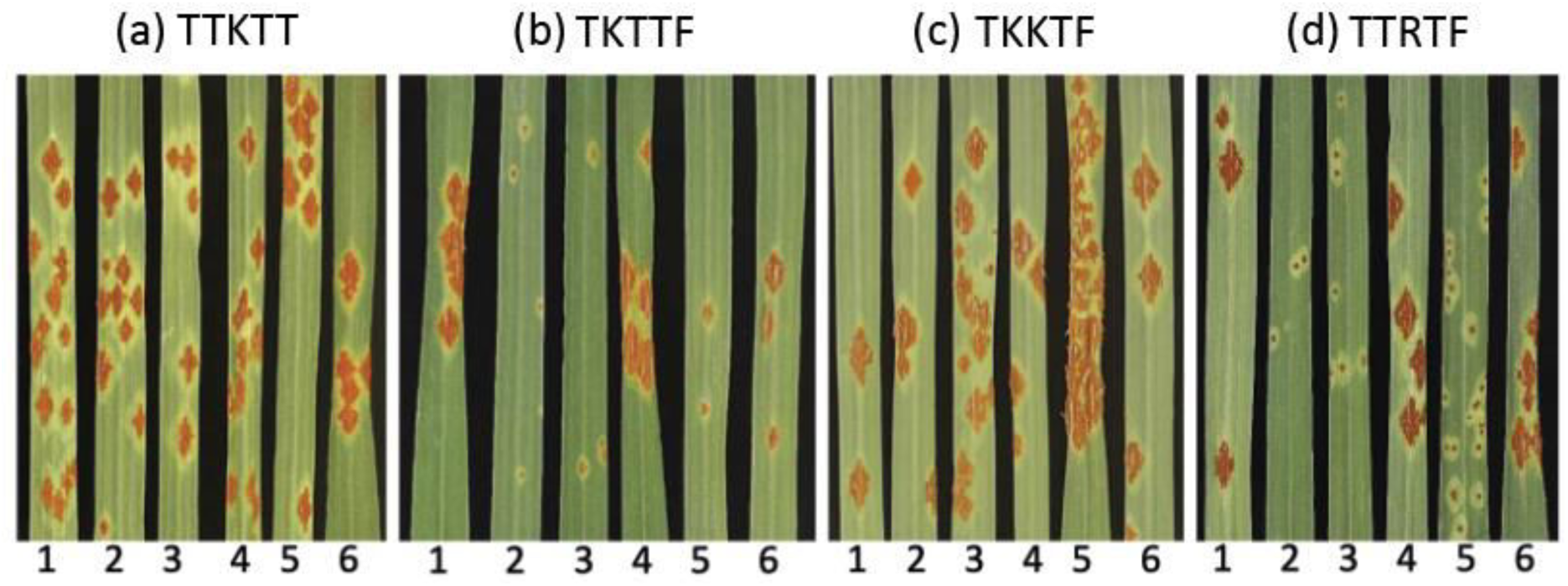
The *Ae. tauschii* stem rust resistance gene *SrTA1662* maintains race specificity as a transgene in wheat. The *SrTA1662* gene was transformed into the stem rust susceptible wheat cultivar Fielder. Shown are T_2_ generation lines selected to be homozygous for the transgene or to be non-transgenic segregants. **a**, Inoculation with isolate IT200a/18 (race TKKTF). **b**, Inoculation with isolate IT16a/18 (race TTRTF). **c**, Inoculation with isolate ET11a/18 (TKTTF). **d**, Inoculation with isolate KE184a/18 (Kenya). Numbering refers to 1 = DPRM0050 (null of DPRM0051), 2 = DPRM0051, 3 = DPRM0059, 4 = DPRM0062 (null of DPRM0059), 5 = DPRM0071, 6 = DPRM0072 (= null of DPRM0071) (see Supplementary Table E).

